# A molecular mechanism for the procentriole recruitment of Ana2

**DOI:** 10.1101/647149

**Authors:** Tiffany A. McLamarrah, Sarah K. Speed, Daniel W. Buster, Carey J. Fagerstrom, Brian J. Galletta, Nasser M. Rusan, Gregory C. Rogers

**Affiliations:** Department of Cellular and Molecular Medicine, University of Arizona Cancer Center, University of Arizona, Tucson, AZ 85724 USA; National Heart, Lung, and Blood Institute (NHLBI), National Institutes of Health, Bethesda, MD 20892 USA

**Author notes:** Correspondence to Nasser M. Rusan; or Gregory C. Rogers. N.M. Rusan and G.C. Rogers contributed equally to this paper.

## Abstract

Centriole duplication begins with the assembly of a pre-procentriole at a single site on a mother centriole and proceeds with the hierarchical recruitment of a conserved set of proteins, including Polo-like kinase 4 (Plk4)/ZYG-1, Ana2/SAS-5/STIL, and the cartwheel protein Sas6. During assembly, Ana2/STIL stimulates Plk4 kinase activity, and in turn, Ana2/STIL’s C-terminus is phosphorylated, allowing it to bind and recruit Sas6. The assembly steps immediately preceding Sas6-loading appear clear, but the mechanism underlying the upstream pre-procentriole recruitment of Ana2/STIL is not. In contrast to proposed models of Ana2/STIL recruitment, we recently showed that *Drosophila* Ana2 targets procentrioles independent of Plk4-binding. Instead, Ana2 recruitment requires Plk4 phosphorylation of Ana2’s N-terminus, but the mechanism explaining this process is unknown. Here, we show that the amyloid-like domain of Sas4, a centriole surface protein, binds Plk4 and Ana2, and facilitates phosphorylation of Ana2’s N-terminus which increases Ana2’s affinity for Sas4. Consequently, Ana2 accumulates at the procentriole to induce daughter centriole assembly.

## Introduction

Centrosomes serve as microtubule-nucleating and organizing centers within cells (Conduit et al., 2015). At the core of these organelles lie barrel-shaped centrioles composed of a radial array of nine microtubules (or microtubule bundles) along with several interconnecting proteins (Ito and Bettencourt-Dias, 2018). Centrioles function to recruit a cloud of pericentriolar material (PCM) which nucleates microtubule growth (Mennella et al., 2014). Position mapping of centrosomal proteins has revealed three hierarchal zones within the organelle that emanate from the centriole center: the centriole, bridge, and PCM zones (Varadarajan and Rusan, 2018). Proteins residing in the bridge zone, such as Sas4/CPAP and Asl/Cep152, are embedded in and/or extend away from the centriole surface and act as scaffolds to recruit and anchor PCM proteins (Fu and Glover, 2012; Lawo et al., 2012; Mennella et al., 2012; Sonnen et al., 2012). Importantly, bridge proteins also play a crucial role in centrosome duplication (Banterle and Gönczy, 2017).

Centrioles are the duplicating elements of centrosomes, a process that is regulated in a cell cycle-dependent manner (Nigg and Holland, 2018). Normally, G1-phase cells contain two centrioles that each spawn an orthogonally-positioned ‘daughter’ (also known as a procentriole) during S-phase. During mitotic progression, new daughter centrioles then sequentially recruit the final structural components and bridge proteins that convert them into mature structures capable of accumulating PCM, thereby enabling them to function as centrosomes in the next cell cycle (Wang et al., 2011). For example, *Drosophila* Sas4, which is present on daughter centrioles as cells enter mitosis, is essential for the mitotic loading of its binding partner Asl (Dzhindzhev et al., 2010; Novak et al., 2014; Fu et al., 2016).

Although the earliest physical manifestation of procentrioles appears during S-phase (Robbins et al., 1968), duplication in flies begins during mitosis with the recruitment of the master-regulator Polo-like kinase 4 (Plk4). Plk4 activity is necessary for centriole assembly and is sufficient to induce centriole overduplication when the kinase is overexpressed (Bettencourt-Dias et al., 2005; Habedanck et al., 2005; Kleylein-Sohn et al., 2007; Peel et al., 2007; Rodrigues-Martins et al., 2007; Holland et al., 2010). Initially, Plk4 interacts with a centriole-targeting factor, such as Asl (Cep152 in humans) (Cizmecioglu et al., 2010; Dzhindzhev et al., 2010; Hatch et al., 2010; Kim et al., 2013; Sonnen et al., 2013), and, during late mitosis, appears as a single asymmetric spot on each mother centriole, a structure called the ‘pre-procentriole’ from which daughter centrioles assemble (Dzhindzhev et al., 2017). Characterizing the functional consequences of Plk4 phosphorylation of its substrates is key to understanding centriole assembly.

*Drosophila* Anastral Spindle 2 (Ana2; STIL in humans) is an essential centriole zone protein (Goshima et al., 2007), and colocalizes with Plk4 on pre-procentrioles. Ana2 contains an N-terminal (NT) Sas4 binding domain, a central coiled-coil (CC), and a C-terminal (CT) STil/ANa2 (STAN) domain (Fig. 1 A) (Stevens et al., 2010; Tang et al., 2011; Vulprecht et al., 2012; Cottee et al., 2013; Hatzopoulos et al., 2013). Through interactions with its CC and CT, Ana2/STIL binds Plk4 (Dzhindzhev et al., 2014; Ohta et al., 2014; Kratz et al., 2015; McLamarrah et al., 2018; Ohta et al., 2018), and activates the kinase, possibly by relieving Plk4 autoinhibition (Klebba et al., 2015; Arquint et al., 2015; Moyer et al., 2015). In turn, Ana2/STIL is extensively phosphorylated (Dzhindzhev et al., 2014; Ohta et al., 2014; Kratz et al., 2015), which occurs in an ordered pattern (Fig. 1 A) (McLamarrah et al., 2018). Initially, phosphorylation predominantly occurs in the NT, which promotes Ana2 recruitment to the procentriole assembly site (Dzhindzhev et al., 2017). Second, phosphorylation of the STAN domain generates a phospho-binding site for the cartwheel protein, Sas6, a critical step required for Sas6 procentriole loading (Dzhindzhev et al., 2014; Ohta et al., 2014; Kratz et al., 2015; Moyer et al., 2015). Lastly, phosphorylation of the Ana2 central region releases Plk4 (McLamarrah et al., 2018).

**Figure 1.**
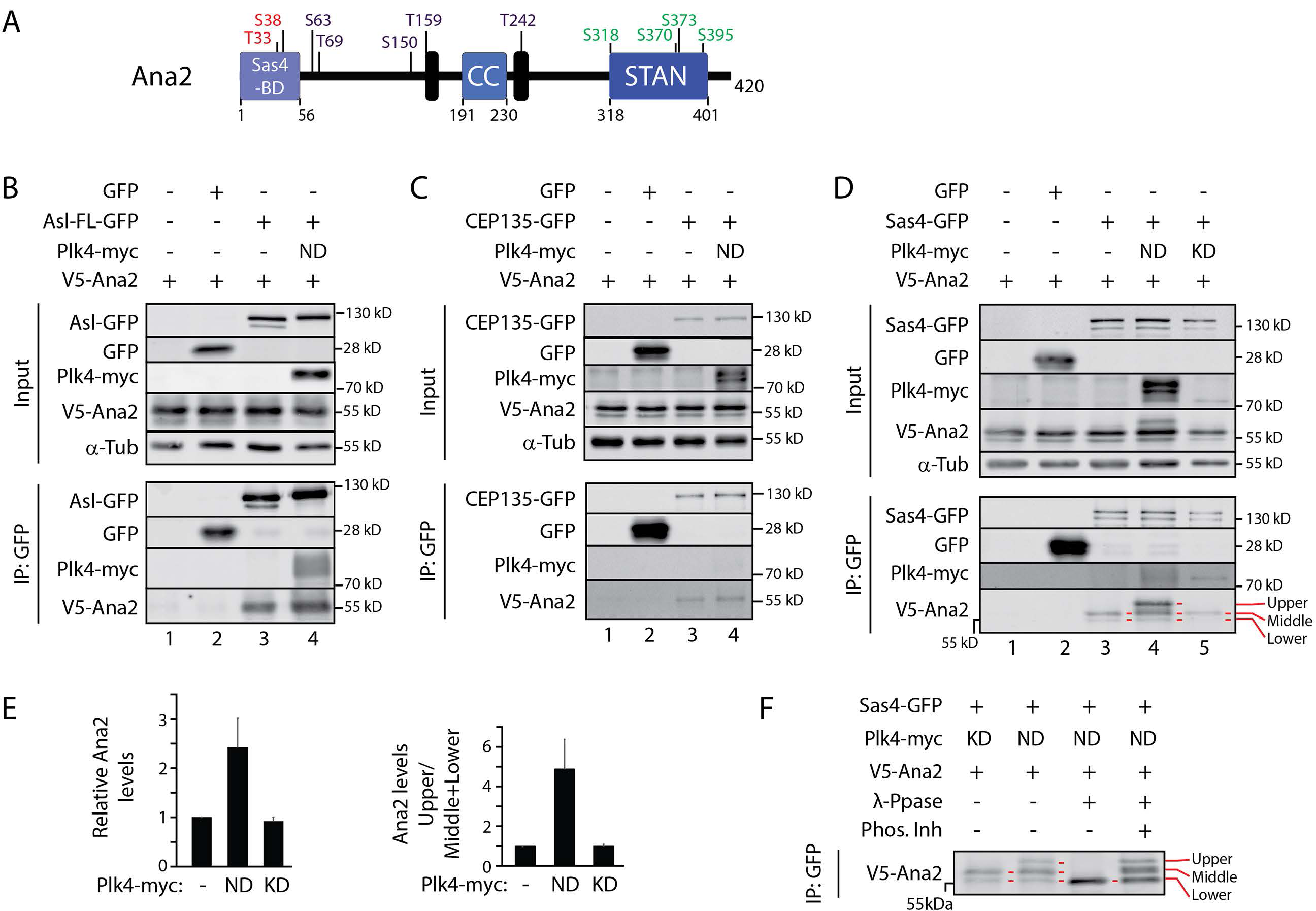
Plk4 activity enhances the interaction between Ana2 and Sas4, producing a hyperphosphorylated Ana2 species. **(A)** Linear map of Ana2 depicting functional domains and Plk4 phospho-sites. Ana2 is phosphorylated by Plk4 in an ordered pattern, first on NT residues T33/S38 (red) followed by STAN domain residues S318/S370/S373/S395 (green), and then central residues S63/T69/S150/T159/T242 (purple) (McLamarrah et al., 2018). Sas4-binding domain (SBD) (Cottee et al., 2013), LC8 binding-sites (black boxes) (Slevin et al., 2014), and coiled coil (CC) are shown. **(B-D)** Ana2 binds Cep135 (**B**), Asl (**C**) and Sas4 (**D**), but co-expression of catalytically-active non-degradable (ND) Plk4 enhances just the association with Sas4. S2 cells were co-transfected with the indicated constructs and the next day induced to express for 24 hours by the addition of 1mM CuSO_4_. Anti-GFP IPs were then prepared from lysates, and Western blots of the inputs and IPs probed for GFP, V5, myc, and α-tubulin. Note, Ana2 presents as three different phospho-species (red lines in all figures) indicated as lower (non-phospho), middle (phospho), and upper (hyperphospho) species. Co-expression of active Plk4 (but not kinase-dead) and Sas4 generates a pool of hyperphospho-Ana2 (**D**). **(E)** Graphs show relative Ana2 intensities in the IPs shown in **D**, lanes 3-5. (Left panel) Ana2 intensities within a region of interest containing all Ana2 species were measured and then normalized to the treatment lacking Plk4 expression. (Right panel) Graph shows ratio of the upper (hyperphosphorylated) to the middle and lower Ana2 species for the three indicated treatments. n = 3 experiments. Error bars, SEM. **(F)** Co-expression of active Plk4 and Sas4 generates a hyperphosphorylated form of Ana2. Anti-GFP IPs were prepared from cell lysates expressing the indicated proteins and then were either mock-treated (lanes 1 and 2), or incubated with λ-phosphatase (λ-Ppase) (lane 3) or λ-Ppase plus phosphatase inhibitor cocktail (Phos Inh).

While the mechanism of Sas6 recruitment to the pre-procentriole appears clear, it is unclear how the immediate upstream step of Ana2 recruitment occurs. Based on the finding that deletion of the central CC in Ana2 prevents Plk4 binding and its targeting to centrioles (Ohta et al., 2014; Arquint et al., 2015; Kratz et al., 2015; Moyer et al., 2015), it has been proposed that a direct interaction with Plk4 mediates Ana2 loading onto centrioles. However, we have shown that CC deletion actually has multiple deleterious effects because it also disrupts Ana2’s ability to oligomerize and interact with Sas4 (McLamarrah et al., 2018). By using a separation-of-function phosphomimetic mutation in Ana2 that is unable to bind Plk4, we observed that phosphomimetic Ana2 localizes properly to the procentriole assembly site (McLamarrah et al., 2018), suggesting that Ana2 recruitment relies on a mechanism independent of Plk4 binding. Importantly, though, Ana2/STIL localization to centrioles requires Plk4 kinase activity (Moyer et al., 2015; Dzhindzhev et al., 2017), consistent with the observation that phosphorylation of the Ana2 NT is necessary for procentriole recruitment. How Ana2 phosphorylation promotes its targeting to the procentriole assembly site is unknown. Here, we provide the first molecular mechanism of Ana2 recruitment, a multiple step process coordinated through Sas4 to achieve Ana2 enrichment on the procentriole assembly site.

## Results

### Sas4 promotes Ana2 hyperphosphorylation by Plk4

*Drosophila* Ana2 does not require direct binding to Plk4 for its recruitment to procentrioles (McLamarrah et al., 2018). However, centriole recruitment of Ana2/STIL does require Plk4 kinase activity as well as phosphorylation of its N-terminus (NT) (Moyer et al., 2015; Dzhindzhev et al., 2017). Therefore, we hypothesized that Plk4 activity enhances the interaction between Ana2 and a protein that resides at or near the centriole surface. Cep135, Asl and Sas4 are excellent candidates because they fulfill this spatial criterium and are also known to directly bind Ana2 (Cottee et al., 2013; Hatzopoulos et al., 2013; Galletta et al., 2016). To test whether Plk4 activity increases the association of Ana2 with these proteins, we co-expressed a highly-active, non-degradable (ND) form of Plk4-myc with V5-Ana2 in S2 cells along with GFP-tagged Asl, Cep135 or Sas4. We then performed GFP immunoprecipitations (IP) and examined the amount of V5-Ana2 in the immunoprecipitate. As expected, Ana2 associated with all three proteins without Plk4 overexpression, but not control GFP (Fig. 1, B, C and D, lanes 1-3). In cells co-expressing Plk4, we found that Ana2 levels remained unchanged in Asl and Cep135 IPs (Fig. 1, B and C, lanes 4). However, Plk4 promoted a ∼2.5-fold increase in the amount of Ana2 that associated with Sas4 (Fig. 1, D, lane 4; E, left panel). Notably, this effect required Plk4 kinase activity because Ana2 enrichment was not observed from cells co-expressing kinase-dead (KD) Plk4 (Fig. 1 D, lane 5; E, left panel).

Closer examination of the immunoblots revealed that Ana2 normally exists as a tight doublet on SDS-PAGE (Fig. 1, D, lane 3), suggesting that Ana2 exists in a phosphorylated state. Strikingly, Sas4 and Plk4 co-expression shifted Ana2 to an even lower electrophoretic mobility (Fig. 1, D, lane 4, upper band). This shift in Ana2 mobility was not observed with co-expression of Plk4 and Asl or Cep135 (Fig. 1, B and C), nor was it seen with the combined expression of Sas4 and kinase-dead Plk4 (Fig. 1, B-D), indicating that co-expression of Sas4 and Plk4 leads to a new hyperphosphorylated Ana2 species. This was confirmed by phosphatase treatment of the IPs which collapsed both middle and upper Ana2 isoforms to the lower non-phosphorylated species; the effect of the phosphatase on collapsing the Ana2 triplet was prevented by addition of phosphatase inhibitor (Fig. 1 F). Moreover, we measured a 4-fold increase in the amount of hyperphosphorylated Ana2 that bound Sas4 when Plk4 was catalytically active compared to cells expressing inactive Plk4 (Fig. 1 E, right panel). Taken together, our findings suggest that Plk4 activity enhances the interaction between Sas4 and Ana2.

### Ana2 phosphorylation by Plk4 requires interaction between Ana2 and the Sas4 G-box

We next asked which domain in Sas4 is responsible for the increased binding and production of hyperphosphorylated Ana2 in cells overexpressing Plk4. Sas4 is a multidomain protein containing an N-terminal PN2-3 domain, an adjacent microtubule-binding domain, a central coiled-coil, and ends with the amyloid-like G-box/TCP domain (Hung et al., 2004; Hsu et al., 2008; Hatzopoulos et al., 2013; Zheng et al., 2014; Cutts et al., 2015; Sharma et al., 2016; Zheng et al., 2016). We first carved Sas4 into three domains named A, B and C (Fig. 2 A), and expressed each as a GFP-fusion along with V5-Ana2 and Plk4-myc (either active or kinase-dead) in S2 cells. As before, GFP IPs were performed to analyze for differences in Ana2 binding and mobility on SDS-PAGE. We found that only the G-box of Sas4 was required for Ana2 binding (Fig. 2 B, lanes 3, 4, 9-12). In addition, Ana2 bound the G-box regardless of expressed Plk4 activity (Fig. 2 B, lanes 3 vs 4 and 9 vs 10). These results confirm previous work that found Ana2/STIL NT bound the G-box regardless of phospho-state (Cottee et al., 2013; Hatzopoulos et al., 2013). Notably, expression of active Plk4 did have two effects. First, active Plk4 increased the level of hyperphosphorylated Ana2 (Fig. 2 B, inputs, lanes 3 vs 4 and 9 vs 10), and, second, active Plk4 increased the amount of bound Ana2 (Fig. 2 B, IPs, lanes 3 vs 4 and 9 vs 10). Expression of a truncated Sas4 mutant lacking the G-box (ΔC) failed to bind Ana2 (Fig. 2 B, lanes 11 and 12). Thus, within the context of this assay, our findings suggest that the Sas4 G-box by itself is necessary and sufficient for Plk4 to hyperphosphorylate Ana2 and enhance the Ana2-Sas4 interaction.

**Figure 2.**
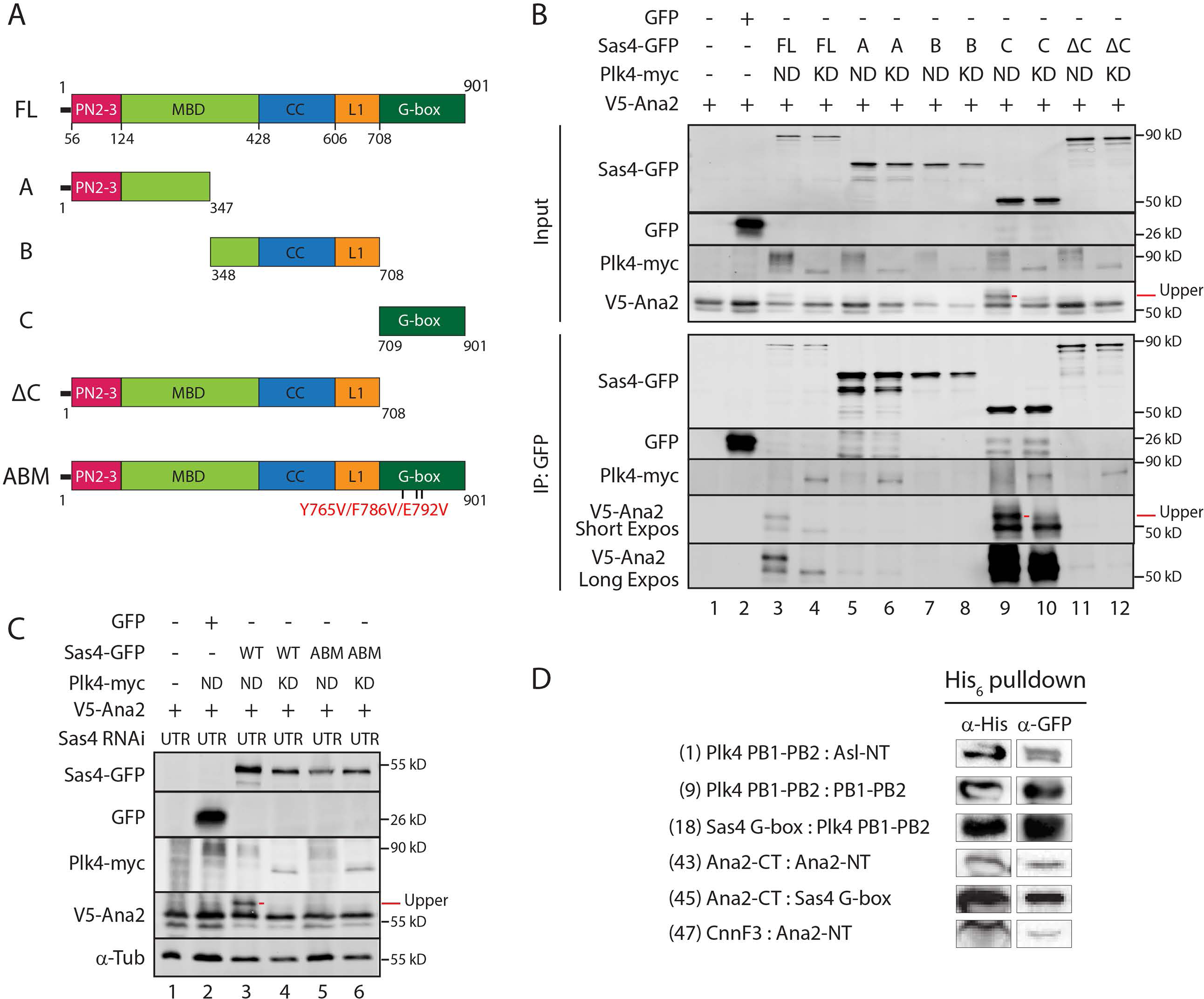
Binding between Ana2 and the Sas4 G-box is required for Plk4-dependent phosphorylation of Ana2 and for enhanced Ana2-Sas4 association. **(A)** Linear maps of Sas4 constructs used in these experiments that depict functional and structural domains. MBD, microtubule-binding domain; CC, coiled-coil; linker 1, L1. The Ana2-binding mutant (ABM) harbors the indicated amino acid substitution (Bond et al., 2005; Cottee al et., 2013). **B** Sas4 G-box promotes hyperphosphorylation of Ana2 by Plk4. S2 cells were co-transfected with the indicated constructs and the next day induced to express for 24 hours by the addition of 1mM CuSO_4_. Anti-GFP IPs were then prepared from lysates, and Western blots of the inputs and IPs probed for GFP, V5, and myc. Hyperphosphorylated Ana2 isoform is clearly diminished in the presence of kinase-dead (KD) Plk4 or absence of the Sas4 G-box. **C** An Ana2-binding mutation (ABM) in the Sas4 G-box disrupts Ana2 hyperphosphorylation by Plk4. S2 cells were depleted of Sas4 by RNAi targeting its untranslated region (UTR) for 6 d. On day 4, cells were transfected with the indicated constructs and induced to express the next day for 24 h by the addition of 1mM CuSO_4_. Proteins were detected by immunoblotting cell lysates with anti-GFP, V5, myc and α-tubulin. Note that hyperphosphorylated Ana2 is absent when endogenous Sas4 is replaced with Sas4-ABM. **D** 6 protein complexes were identified using the pExpress-Dual screen. Anti-His immunoprecipitates were probed with anti-His and GFP. IPs Interaction numbers are indicated (see Fig. S4 B).

Previous work revealed that the Sas4-Ana2 interaction is highly conserved, and structural studies of the complex show direct binding between a short segment of the Ana2 NT and the single β-sheet comprising the G-box (Cottee et al., 2013; Hatzopoulos et al., 2013). This interaction is clearly important: G-box mutations that disrupt Ana2 binding prevent centriole duplication and are linked to primary microcephaly (Thornton and Woods, 2009; Cottee et al., 2013). Although a structural role within the cartwheel has been proposed for the Ana2-G-box interaction, at present it is unknown how this interaction promotes centriole assembly at a mechanistic level. Therefore, we asked whether disrupting the Ana2-G-box interaction could prevent Plk4 from phosphorylating Ana2. Based on the atomic structure of the fly complex (Cottee et al., 2013), we made substitutions in three key G-box residues (Y765V/F786V/E792V) designed to specifically abolish Ana2 binding, which included the microcephaly equivalent mutation E792V (E1235V in human Sas4) (Bond et al., 2005). We named this the Ana2-Binding Mutant (ABM) (Fig. 2 A). Endogenous Sas4 was depleted from S2 cells using RNAi and then replaced with either GFP, full-length wild-type (WT) Sas4-GFP or Sas4-ABM-GFP. V5-Ana2 was hyperphosphorylated when active Plk4 was co-expressed with WT-Sas4 (Fig. 2 C, lane 3) but not when Sas4 was absent (Fig. 2 C, lane 2) or when the binding mutant, Sas4-ABM, was co-expressed (Fig. 2 C, lane 5). Similarly, endogenous Ana2 was hyperphosphorylated in the presence of active Plk4 when Sas4-WT (but not Sas4-ABM) was expressed (Fig. S1 A). Lastly, expression of the Sas4-C fragment containing the ABM mutations also failed to hyperphosphorylate Ana2 when co-expressed with active Plk4 (Fig. S1 B). Taken together, our findings provide the first mechanistic insight into the conserved Ana2-G-box interaction, namely that it enables Plk4 to phosphorylate Ana2, which then enhances Ana2 binding to Sas4.

### The Sas4 G-Box binds the Plk4 Polo Box 1-2 cassette

How could the Sas4 G-box facilitate Plk4 to phosphorylate Ana2? One possibility is that the G-box, which forms a single extended β-sheet (Cutts et al., 2015), acts as a binding platform for both Plk4 and Ana2, placing them in a proximity and orientation that stimulates Ana2 phosphorylation. However, direct binding between Sas4 and Plk4 has not been shown and was not previously detected in our yeast two-hybrid (Y2H) map of the fly centrosome (Galletta et al., 2016). To identify new interactions between fly centrosome proteins, we developed a high-throughput protein screen for proteins that form stable binary complexes in bacteria. 17 centrosome genes were selected for analysis, including Sas4, Plk4 and Ana2 (Figs. S1 C, D).

Full-length genes and fragments were tagged with either His_6_ or GFP and randomly inserted in a bacterial dual-tag expression plasmid (Fig. S2 A and A’, Table S1). In brief, bacterial lysates were subjected to a two-step affinity-purification scheme to isolate protein complexes. From a possible 992 protein combinations, we identified 90 interactions from a primary screen that involved positive detection of GFP in His_6_-tagged pulldowns (Fig. S2 B, S3). Duplicates and false-positive dual-tagged single proteins were eliminated by sequencing the original colonies, leaving a possible 48 positive interactors. Binding was further tested using a secondary screen consisting of tandem and reciprocal pulldown/IPs (Fig. S2 B, S4). In total, we found 6 high-confidence interactors (Fig. 2 D, S2 B), including binding between 1) the central Plk4 Polo Box (PB) cassette PB1-PB2 with itself, 2) PB1-PB2 and the NT of Asl, and 3) NT and CT fragments of Ana2; all three are well-described interactions that validated our screen (Slevin et al., 2012; Dzhindzhev et al., 2010; McLamarrah et al., 2018). Remarkably, we also identified the G-box as another PB1-PB2 binding partner as well as an interaction between the G-box and a CT fragment of Ana2 lacking the previously described G-box-binding motif. An interaction between the Ana2-NT and Cnn was also detected but is not examined further in this study. Thus, our findings suggest that Plk4 can directly bind the G-box through its PB1-PB2 cassette and that Ana2 may contains additional undescribed G-box binding sites.

### Sas4 promotes Plk4 phosphorylation of Ana2 S38

We next sought to determine which Ana2 residue(s) are primarily phosphorylated in the presence of Sas4. Several potential sites exist, as Ana2 is phosphorylated by Plk4 on at least 11 residues along its length (Fig. 1A) (Dzhindzhev et al., 2014; McLamarrah et al., 2018). It follows that substitution of the relevant Ser/Thr residues with non-phosphorylatable alanine (A) would block the shift in Ana2 mobility when co-expressed with Sas4 and Plk4, whereas phosphomimetic (PM) Asp/Glu substitutions would produce electrophoretic shifts in cells containing inactive Plk4. We first examined whether Sas4 promotes Plk4 phosphorylation of the STAN domain. Although modification of this region is not required for centriole targeting in fly cells (Dzhindzhev et al., 2014), it does control the localization of STIL in cultured human cells (Moyer et al., 2015). Therefore, we made substitutions of the three most conserved phosphorylated STAN residues (S318/S370/S373) (generating nonphosphorylatable ‘3A’ and phosphomimetic ‘3PM’ mutants) and then examined V5-Ana2 mobility by Western blotting Sas4-GFP IP samples. Both Ana2-3A and 3PM co-immunoprecipitated with Sas4 and shifted to slower migrating isoform in cells co-expressing active Plk4 (Fig. S5 A, lanes 5 and 7), demonstrating that STAN domain phosphorylation is not responsible for the Sas4-dependent shift in Ana2 mobility.

We next tested whether Sas4 promotes phosphorylation of Ana2’s NT by examining mutants with substitutions (alanine ‘2A’, or glu/asp ‘2PM’) of both phospho-residues T33 and S38 (Fig. 1 A). We found that, despite its ability to co-IP with Sas4, Ana2-2A failed to shift to the hyperphosphorylated form and its levels did not increase in the immunoprecipitates when co-expressed with active Plk4 (Fig. S5 B, lanes 5). In contrast, Ana2-2PM produced the mobility shift regardless of Plk4’s catalytic state (Fig. S5 B, lanes 7 and 8). Thus, Sas4 increases the Plk4-dependent phosphorylation of Ana2 NT.

To determine whether the Ana2 mobility shift and enhanced binding to Sas4 were due to modification of a single residue, we generated individual T33 and S38 phospho-mutants and, as before, examined their mobilities by Western blotting Sas4-GFP IP samples. We found that only one mutation, S38A, prevented the Ana2 mobility shift and its accumulation in immunoprecipitates when co-expressed with Sas4 and active Plk4 (Fig. S5 C, lanes 9 and 10).

Phosphomimetic S38D displayed the mobility shift regardless of Plk4 activity (Fig. S5 C, lanes 11 and 12). Identical changes in Ana2 mobility were also detected in whole lysates prepared from cells depleted of endogenous Ana2 (Fig. 3 A). Our data indicate that the Ana2-Sas4 interaction is required for Plk4 to phosphorylate Ana2 (Fig. 2 C), and failure of the S38A mutant to shift to the slower migrating species could be explained if S38 phosphorylation were required for Sas4 binding. Because Ana2 oligomerizes (Cottee et al., 2015), we again used RNAi to eliminate the influence of endogenous Ana2 on the IP assay. Interestingly, in each case, S38 phospho-mutants were still able to bind Sas4 regardless of Plk4 activity (Fig. 3 B, lanes 6-9). Thus, our findings indicate that Sas4 directs Plk4 to phosphorylate S38 of Ana2, although S38 phosphorylation is not required for Ana2 to bind Sas4.

**Figure 3.**
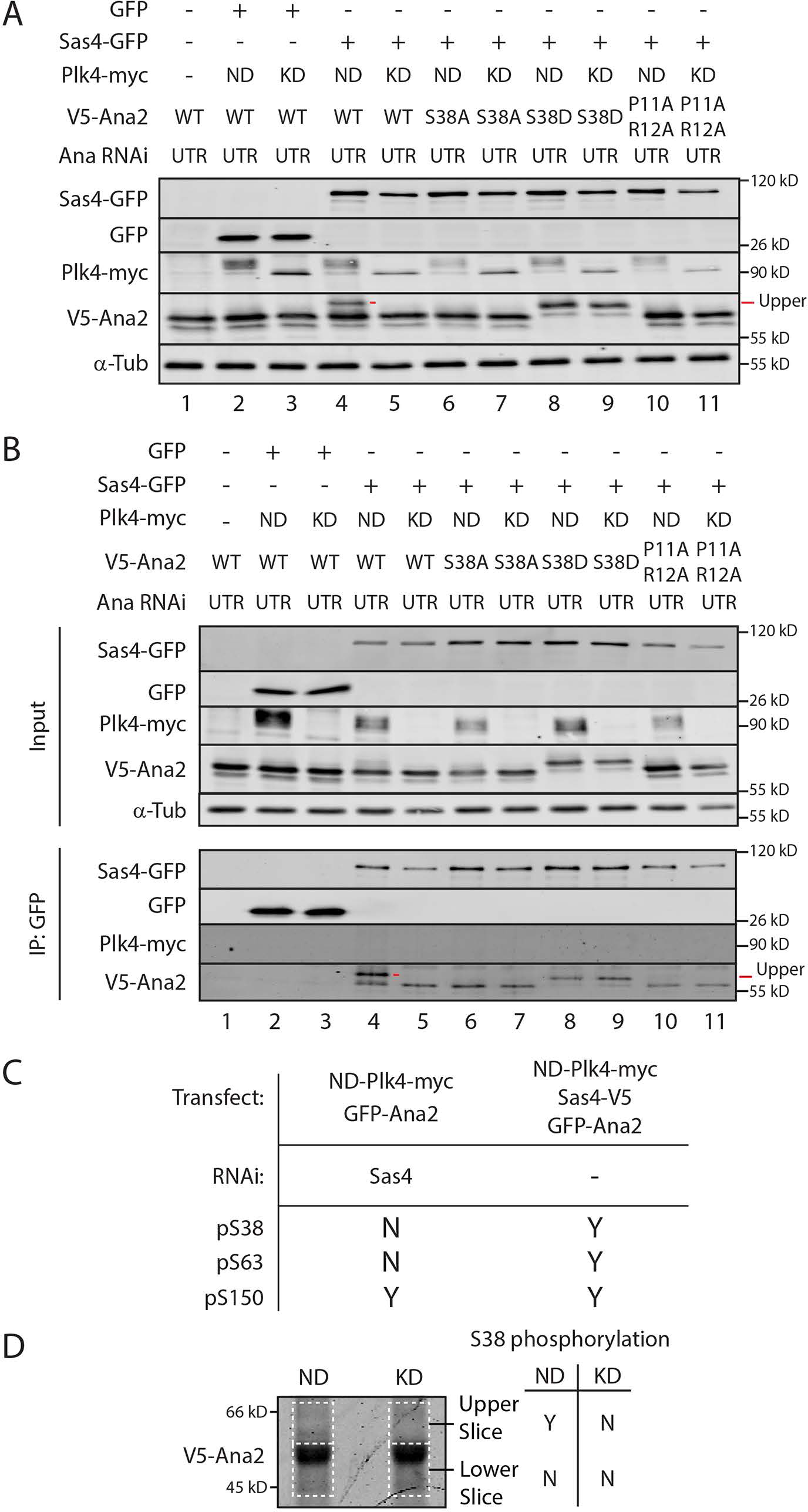
Sas4 stimulates Plk4 phosphorylation of Ana2 S38. **(A and B)** The phospho-null mutant S38A blocks the shift in Ana2’s electrophoretic mobility, as does the Sas4-binding mutation (SBM) P11A/R12A. S2 cells were Ana2 depleted by RNAi targeting the untranslated region (UTR) for 6 d. On day 4, cells were transfected with the indicated constructs and induced to express the next day for 24 h by the addition of 1mM CuSO_4_. Whole cell lysates (**A**) or anti-GFP IPs were then prepared (**B**) and Western blots of the samples probed for GFP, V5, myc, and α-tubulin. **C** Table shows the presence (Yes or No) of N-terminal phospho-residues in Ana2 IPed from cells with the indicated treatments. **D** Coomassie-stained protein gel of immunoprecipitated Ana2 from S2 cells co-expressing Sas4-GFP and either active (ND) or kinase-dead (KD) Plk4-myc. Gel slices corresponding to the upper and lower migratory species of Ana2 (dashed lines) were isolated and prepared for MS/MS analysis. Detection (Yes or No) of phospho-S38 within each gel region by MS/MS is shown.

If Ana2-Sas4 binding is required for Ana2 hyperphosphorylation, then a Sas4-binding mutation in Ana2 should also prevent its hyperphosphorylation when co-expressed with Sas4 and Plk4. To test this, we depleted cells of endogenous Ana2 and replaced it with a P11A/R12A double mutant; P11 and R12 lie in the conserved PRxxPxP motif that directly interacts with the G-box based on atomic structure of the complex (Cottee et al., 2013; Hatzopoulos et al., 2013). Although the double mutation was not shown to disrupt Sas4 binding, analogous mutations in zebrafish STIL dramatically decrease Sas4 binding (Cottee et al., 2013). Nevertheless, Ana2-P11A/R12A does not fully rescue centrosome duplication in Ana2 mutant embryos (Cottee et al., 2013). We found that Ana2-P11A/R12A was able to co-IP with Sas4 from Ana2-depleted cell lysates but at reduced levels compared to WT-Ana2 (Fig. 3 B, lane 10 and 11). Ana2-P11A/R12A also failed to shift to the hyperphosphorylated form as assayed by Western blotting of whole cell lysates and Sas4-GFP IPs (Fig. 3 A and B, lanes 10 and 11). Thus, our results indicate that an intact PRxxPxP motif is required for Ana2 phosphorylation by Plk4.

Our previous in vitro kinase assays and tandem mass spectrometry (MS/MS) of Ana2 revealed that S38 is one of the first residues phosphorylated by Plk4 (McLamarrah et al., 2018). Curiously however, we did not detect S38 phosphorylation when Ana2 was purified from cells overexpressing Plk4. Possibly, this is because Sas4 was not co-overexpressed in these cells, whereas in vitro reactions containing high concentrations of Plk4 and Ana2 can overcome the need for Sas4 to facilitate S38 phosphorylation. To explore this hypothesis, we performed MS/MS analysis of Ana2 immuno-purified from cells co-expressing Plk4 and either depleted of Sas4 or co-expressing Sas4 (Table S2). Phospho-S38 was not detected in Sas4-depleted cells, although phosphorylation of S150, a different Plk4-targeted residue (Dzhindzhev et al., 2014; McLamarrah et al., 2018), was detected (Fig. 3 C). In contrast, expression of Sas4 resulted in detectable phosphorylated S38, S63, and S150 (Fig. 3 C). We next determined which electrophoretic species of Ana2 contained phospho-S38. Ana2 was immuno-purified from cells expressing Sas4 and either catalytically-active (ND) or kinase-dead (KD) Plk4, the IP samples resolved by SDS-PAGE, and then regions from the stained gel separated into upper and lower sections which were analyzed separately by MS/MS (Fig. 3 D, dashed lines). Notably, phospho-S38 was only detected in cells co-expressing Sas4 and ND-Plk4 and only in the upper gel slice where hyperphosphorylated Ana2 migrates. Taken together, our results indicate that Sas4 directs Plk4 to phosphorylate S38 in Ana2, thereby generating the slower migrating Ana2 species.

### Phosphorylation of S38 stimulates Ana2 accumulation on procentrioles

In *Drosophila* cells, Ana2 centriolar recruitment occurs during late anaphase when mother-daughter pairs disengage, typically appearing as two spots that co-localize with each PLP-labeled centriole ring: one spot positioned near the center of the PLP-ring (presumably associated with the cartwheel) and a second spot near the periphery that marks the pre-procentriole. Sas6 follows Ana2 to the pre-procentriole and this pattern persists into interphase (Dzhindzhev et al., 2017). To examine the effects of S38 phospho-mutants on Ana2’s centriole recruitment, we depleted endogenous Ana2 and used super-resolution (SR-SIM) microscopy to localize transgenic GFP-Ana2 in interphase cells. Most centrioles in cells expressing transgenic Ana2-WT contained two Ana2 spots (75%); the remaining centrioles contained only the peripheral procentriole-spot (Fig. 4, rows 1 and 5). Interestingly, linescans of centrioles with two spots revealed an asymmetry in Ana2 levels. Specifically, the immunofluorescence intensity of Ana2 at the procentriole was, on average, 2-fold greater than the intensity of the Ana2 spot located within the mother centriole (Fig. 4, row 1). A similar pattern was observed in cells expressing phosphomimetic Ana2-S38D: 68% of centrioles contained 2 spots of Ana2-S38D where the Ana2 spot on the procentriole was more intense (Fig. 4, rows 3, 7 and 10). However, expression of phospho-null Ana2-S38A altered this pattern. Although some centrioles completely lacked Ana2-S38A (25%), centrioles usually contained the central mother spot but lacked detectable Ana2-S38A on procentrioles (46%) (Fig. 4, rows 6 and 9). Furthermore, linescans of centrioles containing two spots (29%) showed that Ana2-S38A levels were nearly equal at both sites (Fig. 4, row 2). A similar pattern was observed in cells expressing the P11A/R12A mutant, which weakens Sas4 binding and prevents its hyperphosphorylation (Fig. 3, A and B). Only 28% of centrioles contained two spots of Ana2-P11A/R12A and their intensities were generally similar (Fig. 4, row 4). Based on our findings, we suggest that Ana2 loading may occur in stages.

**Figure 4.**
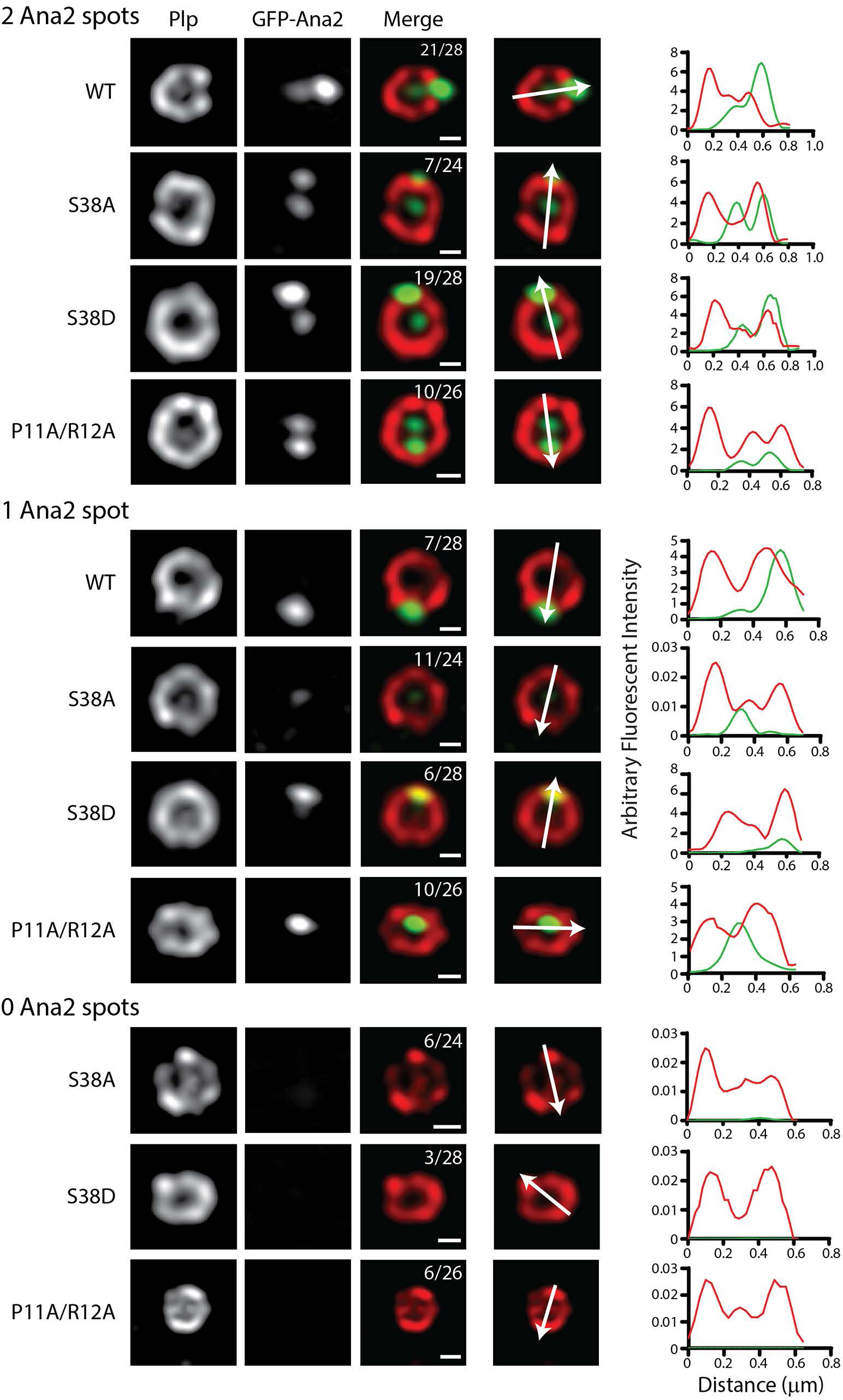
WT and phosphomimetic (S38D) Ana2 accumulate on procentrioles but the phospho-null Ana2-S38A mutant does not. Interphase centrioles in transgenic GFP-Ana2-expressing cells were imaged using super-resolution microscopy. S2 cells were depleted of endogenous Ana2 for 12 days by transfection of UTR dsRNA on days 0, 4 and 8. On days 4 and 8, cells were co-transfected with the indicated GFP-Ana2 constructs (green) and induced to express on day 5 by addition of 0.1 mM CuSO_4_. On day 12, cells were immunostained with anti-PLP (red) to mark the surface of mature centrioles. DNA was visualized with Hoechst (not depicted). Images are grouped as centrioles possessing either two (top), one (middle), or no (bottom) Ana2 spots. Bars, 200 nm. (Right) Graphs of PLP (red) and GFP-Ana2 (green) fluorescence intensities plotted against the length of a linescan (white arrow) spanning the diameter of an individual centriole.

Initially, Ana2 interacts with Sas4 to accumulate at pre-procentrioles at a relatively low basal level (as seen in the S38A and P11A/R12A mutants), but following phosphorylation of S38 (as mimicked by the S38D mutant), Ana2 then accumulates on procentrioles at a several-fold greater level. Failure to accumulate on procentrioles likely explains why the S38A and P11A/R12A mutants fail to rescue centriole duplication defects as we and others previously reported (Cottee et al., 2013; Dzhindzhev et al., 2017; McLamarrah et al., 2018).

### S38 phosphorylation increases Ana2 binding to the Sas4 G-box

Previous studies found human and zebrafish CPAP G-box bound to an N-terminal ∼45 amino acids segment of STIL with a K_D_ of 0.6-2 μM (Cottee et al., 2013; Hatzopoulos et al., 2013). To examine the effects of Ana2 phosphorylation on Sas4 binding, we purified *Drosophila* Ana2 and Sas4 proteins from bacteria and used bio-layer interferometry to measure their binding kinetics (Fig. S6 A-D). We found that an N-terminal fragment (amino acids 1-60) of Ana2 bound the G-box with a K_D_ of 191 nM (Fig. 5 A, S6 B, D), but binding affinity increased over 4-fold (K_D_ = 43 nM) when Ana2 1-60 was pre-phosphorylated with purified Plk4 (Fig. 5 A; S6 B, D).

**Figure 5.**
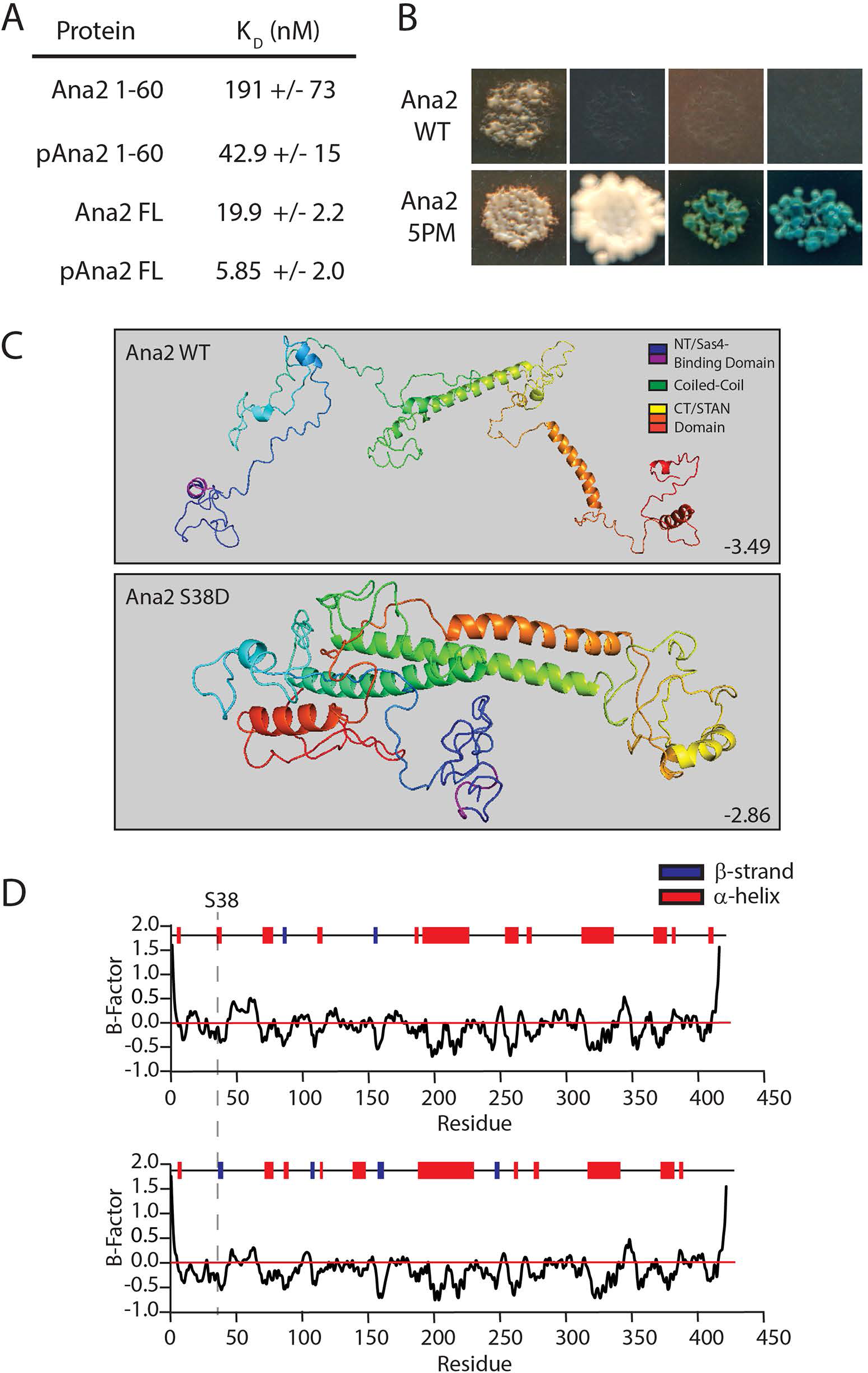
Phosphorylation of Ana2 promotes stronger binding to Sas4. **(A)** Phosphorylated full-length Ana2 displays the tightest binding to the G-Box. Table shows dissociation constants from bio-layer interferometry of Ana2 (untreated or pre-phosphorylated by Plk4) with the Sas4 G-box (amino acids 606-901) using Octet RED384. (See Fig. S6 D.) **B** Phosphomimetic (5PM) full-length (FL) Ana2 shows a strong interaction with FL Sas4 by Y2H analysis. Wild-type (WT) and 5PM Ana2 were screened against Sas4. In each image, colonies from replica plating are shown, and growth indicates the presence of bound bait and prey. Column (1) no selection; (2) growth selection on QDO; growth and color selection on (3) DDOXA and (4) QDOXA (blue color indicates an interaction). **C** I-TASSER protein structure predictions of wild-type (top) and S38D (bottom) Ana2. C-scores are shown. Note that the single S38D substitution is predicted to dramatically alter Ana2 conformation, folding Ana2 so that the N- and CTs are closely positioned. **D** I-TASSER predictions of secondary structure within Ana2. The position of S38 (dotted line) is indicated in a short segment that changes from a α-helix to a β-strand.

Interestingly, full-length Ana2 bound Sas4 with a greater affinity (K_D_ = 20 nM) than the phospho-Ana2 1-60 fragment (Fig. 5 A, S6 C, D), again suggesting that additional Sas4-binding sites may be located outside the NT. When full-length Ana2 protein was pre-phosphorylated by Plk4, it displayed the tightest binding (K_D_ = 6 nM), a ∼3-fold increase compared to the non-phosphorylated protein (Fig. 5 A, S6 C, D). Therefore, for both the NT and FL Ana2 proteins, phosphorylation induced tighter binding to the G-box.

We also inspected the Ana2-Sas4 interaction Y2H. Curiously, no binding was detected between full-length Ana2 and Sas4 (Fig. 5 B). Possibly, the interaction of Sas4 with unmodified Ana2 is too weak to detect by Y2H. Therefore, we examined an Ana2 mutant with 5 phosphomimetic (5PM) substitutions, targeting the 2 N-terminal residues (T33E/S38D) and the 3 conserved phosphorylated STAN residues (S318D/S370D/S373D). We chose to analyze the 5PM mutant because 1) these 5 residues are rapidly phosphorylated by Plk4 in vitro, and 2) the N- and C-termini of Ana2 interact in vitro and this binding is enhanced by these five PM substitutions (McLamarrah et al., 2018), and possibly, binding between the two termini may influence G-box binding. Strikingly, we detected a strong interaction between Sas4 and Ana2-5PM (Fig. 5 B), consistent with our in vitro binding experiments showing that full-length phospho-Ana2 displays the strongest interaction (Fig. 5 A).

Lastly, we examined how phosphorylation of Ana2 S38 could increase binding affinity to Sas4. As suggested by our previous experiments, S38 phosphorylation may induce a structural change in Ana2, stabilizing a folded conformation with increased G-box binding affinity, possibly through multiple sites. Indeed, Ana2 exists in a folded conformation within the cartwheel with its N- and C-termini in close proximity (Gartenmann et al., 2017). Therefore, we used Iterative Threading Assembly Refinement (I-TASSER) structure prediction to better understand how phosphorylation might affect Ana2 folding (Zhang, 2008; Roy et al., 2010, Yang et al., 2016). Accordingly, Ana2 is largely disordered but contains five prominent α-helices of varying length (Fig. 5 C, top). However, a prediction model of Ana2 containing S38D shows a much different folded structure, with the N- and C-termini positioned near each other (Fig. 5 C, bottom). Additionally, S38 lies within a 4 amino acid stretch (V37-L40) that is predicted to be an α-helix in the wild-type sequence but becomes a β-strand when containing the S38D substitution (Fig. 5 D). Remarkably, modeling of NT-2PM (T33E/S38D), CT-3PM (S318D/S370D/S373D) and 5PM Ana2 mutants all predict a folded conformation with β-strand conversion of V37-L40 (Fig. S7). Since Ana2/STIL is a CDK target (Campaner et al., 2005; Zitouni et al., 2016), we also modeled an Ana2 mutant containing Ser/Thr to Asp/Glu substitutions in all 12 of its possible CDK consensus sites (12PM-CDK), but this phosphomimetic mutant did not predict a folded conformation (Fig. S7). Thus, our findings indicate that phosphorylation of the NT and STAN domain promote both Ana2 folding and an interaction between the N- and C-termini, as well as alterations in secondary structure. These structural changes may account for the tighter binding of Ana2 with the Sas4 G-box.

## Discussion

The cartwheel protein Sas6 follows Ana2 to the procentriole assembly site. This event is prompted when Plk4 phosphorylates the Ana2/STIL STAN domain, generating a Sas6-binding surface that supports Sas6 aggregation at the procentriole assembly site (Dzhindzhev et al., 2014; Ohta et al., 2014; Kratz et al., 2015; Moyer et al., 2015). These seminal discoveries provided molecular insight into Plk4’s role in initiating centriole duplication and characterized one of the earliest steps in the process. The goal of our study was to characterize the step prior, specifically, understanding how Ana2 is recruited to the pre-procentriole, the single spot constructed on the surface of a mother centriole during mitotic exit in *Drosophila* cells (Dzhindzhev et al., 2017). We found that Ana2’s centriolar recruitment is triggered by Ana2’s interaction with Sas4, which leads to Ana2 phosphorylation by Plk4, tighter Sas4 binding, and, consequently, its accumulation on procentriole assembly site.

Plk4 phosphorylation of Ana2 S38 is the key to understanding Ana2’s centriole recruitment. S38 is one of the first Ana2 residues phosphorylated by Plk4 (McLamarrah et al., 2018), and the importance of this modification is underscored by the observation that phospho-null S38A fails to load onto most procentrioles and rescue centriole duplication (Fig. 4) (Dzhindzhev et al., 2017). Though Plk4 can phosphorylate Ana2 S38 in vitro in the absence of Sas4 (presumably due to the high concentrations of the reactants), the situation is more complex in cells, and our observations lead us to propose the following model (Fig. 6).

**Figure 6.**
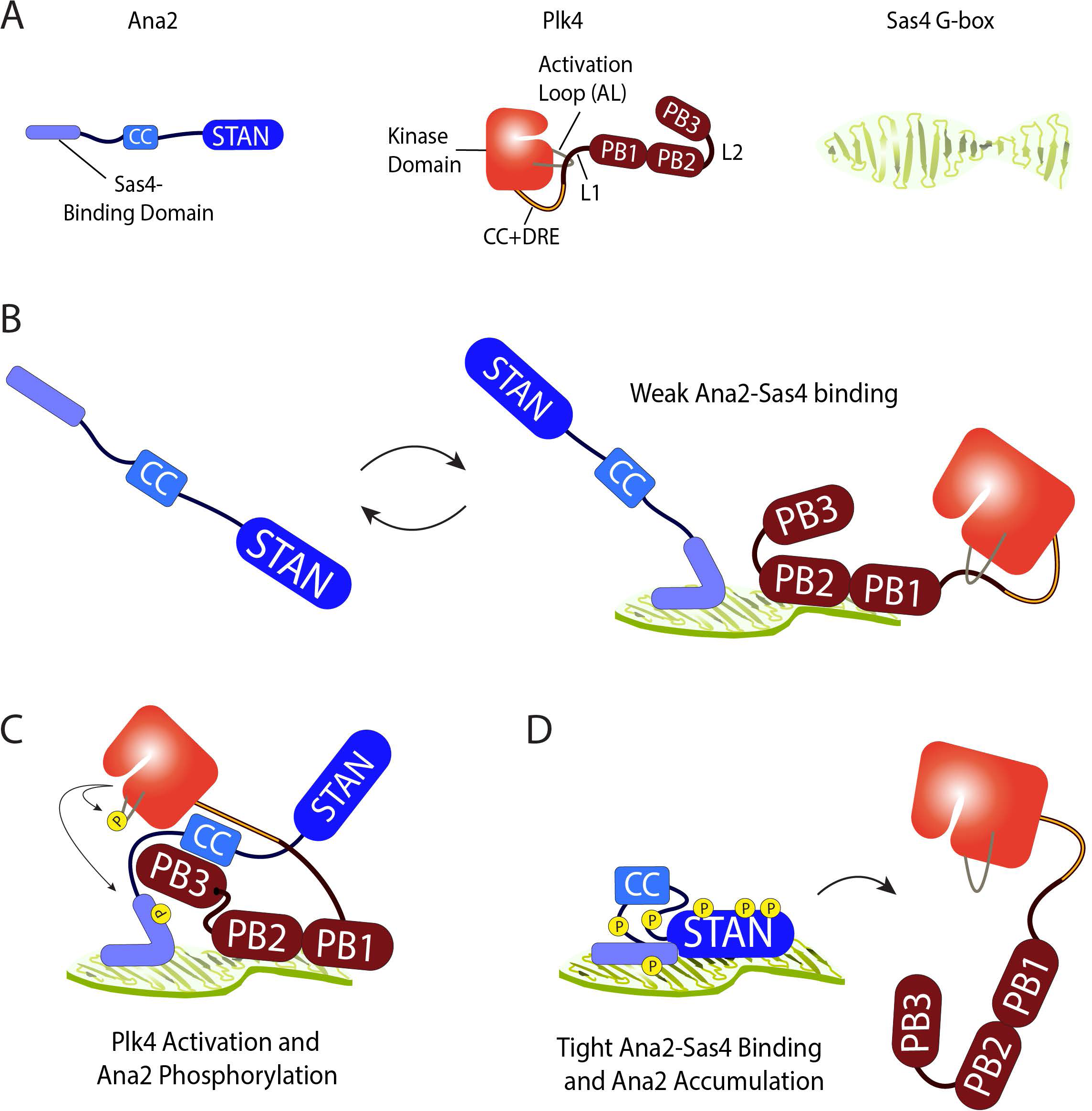
Proposed model depicting the mechanism of Ana2 recruitment to the site of procentriole assembly. **(A)** Schematics of Ana2, Plk4 and Sas4 showing structural and functional domains. Note that Plk4 exists in autoinhibited form. Although these proteins homo-oligomerize, they are shown as monomers for simplicity. Only the C-terminal G-box domain of Sas4 is shown; adopts an extended open β-sheet conformation. **B** During mitotic exit, extended unphosphorylated Ana2 diffusing near a pre-procentriole is weakly bound through its NT to one surface of the Sas4 G-box; substantial exchange of bound Ana2 with the cytoplasmic pool prevents accumulation at the pre-procentriole sufficient to support centriole duplication. Plk4 is also weakly binds the G-box through PB1-PB2. **C** Ana2 binds Plk4 and relieves its autoinhibition. In turn, Plk4 phosphorylates the Ana2 NT, including S38, inducing Ana2 to fold. Consequently, Plk4 phosphorylates the STAN domain. During this step, Plk4 may be transferred from Sas4 to Ana2. **D** Phosphorylated N- and C-terminal domains of Ana2 bind one another and stabilize the folded conformation. Phospho-Ana2 binds tightly to the G-box, through multiple sites of contact. Tight binding to Sas4 allows Ana2 to accumulate on the pre-procentriole, recruit Sas6, and promote centriole duplication. Further phosphorylation of the central domain in Ana2 causes Plk4 to release.

As cells exit mitosis and pre-procentrioles form, Plk4 localizes to this site in an autoinhibited form, utilizing its PB1-PB2 domain to interact with the Sas4 G-box. Although we have not measured in vitro binding parameters of the Plk4-Sas4 interaction, it is likely to be weak because the interaction is undetectable by IP. Ana2 exists as an extended molecule (Fig. 6A) and is first recruited to the centriole surface through a conserved proline-rich motif in its NT that binds the G-box (Cottee et al., 2013; Hatzopoulos et al., 2013). Because the Sas4 G-box binds Ana2 and stimulates the Plk4-dependent hyperphosphorylation of Ana2 (Figs. 1-3), the G-box may act as a multiplex binding platform that juxtaposes Ana2 and Plk4, thereby facilitating the phosphorylation of Ana2 S38 (Fig. 6 B). Future structural studies will be necessary to understand where PB1-PB2 binds the G-box, and if Ana2 and Plk4 can both occupy the G-box simultaneously. (We acknowledge the many possible alternatives; for example, the G-box could be single-occupancy, and incoming Ana2 could displace G-box-bound Plk4, allowing Plk4 to interact with the immobilized Ana2.) Autoinhibited Plk4 is activated by its interaction with Ana2, and, in turn, Ana2 is phosphorylated on its NT (including S38) and then on its STAN domain (Fig. 6 C) (McLamarrah et al., 2018).

S38 is a conserved residue in Ana2/STIL family members and, based on crystal structures of the vertebrate complexes, is in contact with the G-box (Cottee et al., 2013; Hatzopoulos et al., 2013). However, our findings suggest that S38 phosphorylation is structurally transformative, causing a localized change in secondary structure (Fig. 5 C, D) which may promote the stronger G-box binding that we observed (Fig. 5 A). Additional crystallographic studies of phospho- (or phosphomimetic) Ana2 are needed to understand the structural basis of its tighter binding to the G-box; we predict that it does so through more than one G-box binding domain. Further phosphorylation of the N- and C-termini promotes their interaction and, consequently, Ana2 adopts a folded conformation (Fig. 6 D). Tighter Sas4-binding by phospho-Ana2 (Fig. 5 A, B) enhances its accumulation at the future procentriole assembly site compared to phospho-null-Ana2 (Fig. 4). Because the pre-procentriolar accumulation of Ana2 is important for centriole duplication, the apparently unstable binding by phospho-null-Ana2 would explain why this mutant fails to support duplication (Dzhindzhev et al., 2017; McLamarrah et al., 2018). Measuring the turnover of Ana2 and its phospho-isoforms on the pre-procentriole will be a challenging but important line of experimentation.

Lastly, our study identifies the first mechanistic function of the Ana2-G-box interaction. Previous studies demonstrated that this conserved interaction is required for centriole assembly and hypothesized that Ana2-G-box forms a functional ‘strut’ within the pinhead of the cartwheel (Hiraki et al., 2007), thereby providing structural integrity to the organelle (Cottee et al., 2013; Hatzopoulos et al., 2013). Although this hypothesis is plausible, our findings show that this interaction plays a critical step in the assembly process. Without the Ana2-G-box interaction, Plk4 cannot phosphorylate Ana2, likely because Ana2 fails to target pre-procentrioles. In fact, weakening the interaction with the P11A/R12A mutant blocked Ana2 hyperphosphorylation and prevented its normal accumulation on procentrioles. Like Ana2, Sas4 is distributed as an asymmetric spot on centrioles in interphase *Drosophila* cells (Galletta et al., 2016). Determining how and when Sas4 acquires this pattern and whether Plk4 activity regulates its localization will be important future lines of investigation. Identifying additional upstream Plk4 phosphorylation targets is vital in unraveling the complex molecular choreography that underlies centriole assembly.

## Materials and Methods

### Cell culture and double-stranded RNAi

*Drosophila* S2 cell culture and in vitro dsRNA synthesis were performed as previously described (Rogers and Rogers, 2008). Cells were cultured in Sf-900II SFM media (Life Technologies) and RNAi was performed in 6 well plates. To increase the efficiency of RNAi, dsRNA was transfected into S2 cells as described (McLamarrah et al., 2018). For immunoprecipitation assays, 40 μg dsRNA was transfected on day 0 and 20 µg on day 4. For immunofluorescence microscopy and immunoblotting assays, cells were transfected with 40µg dsRNA on days 0, 4, and 8. Control dsRNA and dsRNA targeting Ana2 UTR were generated as previously described (McLamarrah et al., 2018). dsRNA targeting Sas4UTR was synthesized from an oligonucleotide targeting the 5’UTR of Sas4 fused to the 3’UTR of Sas4 to create the following sequence (which includes the T7 RNA polymerase promoter sequence [underlined]): 5’-<u>TAATACGACTCACTATAGG</u>GCGTCTAAACAAAACTGCGATTCTAAGAACGAATATAAAAGCAGTCGGGTCTCTGCTTCCGTTGTTAAGTAAATTAACTTAAGTTTTAATAAATAGCCCTATAGTGAGTCGTATTA-3**’**. Equimolar Sas4UTR oligonucleotides were annealed by first incubating at 95°C for 5 minutes in annealing buffer (10m M Tris, pH 8.0, 50 mM NaCl, 1 mM EDTA) followed by slow cooling to room temperature. In vitro synthesis of Sas4UTR dsRNA was prepared as previously described (McLamarrah et al., 2018) using annealed Sas4UTR oligos as template.

### Immunoblotting

Cells were lysed in PBS-Triton X-100 (0.1%) followed by addition of Laemmli sample buffer. Extracts were boiled for 5 min and stored at −20°C. Samples of equal total protein were resolved by SDS-PAGE, blotted, probed with primary and secondary antibodies, and scanned on an Odyssey imager (Li-Cor Biosciences). For replacement experiments, cells were transfected with dsRNA on days 0, 4, and 8 to eliminate endogenous protein and were concurrently transfected with plasmids encoding protein constructs on days 4 and 8, and then analyzed on day 12. Cells were induced to express with 0.1 mM CuSO_4_ on day 5 and maintained in 0.1 mM CuSO_4_ for the duration of the experiment. Primary antibodies used for Western blotting include rabbit anti-Ana2 (our laboratory), mouse anti-His_6_-HRP conjugated (ThermoFisher), rat anti-Sas4 (our laboratory), mouse anti-V5 monoclonal (Life Technologies), mouse anti-GFP monoclonal JL8 (Clontech), mouse anti-myc (Cell Signaling Technologies), mouse anti-αtubulin monoclonal DM1A (Sigma-Aldrich). Antibody dilutions ranged from 1:500-1:5000. IRDye 800CW secondary antibodies (Li-Cor Biosciences) were prepared according to the manufacturer’s instructions and used at 1:3,000 dilution. Detection was performed using Li-Cor Odyssey CLX or SuperSignal West Dura Extended Duration Substrate and visualized using a ChemiDoc MP Imaging System (Biorad).

### Immunofluorescence microscopy

S2 cells were spread on concavalin A-coated glass bottom plates and fixed in ice cold methanol for 15 minutes. Cells were washed with PBS-Triton X-100 (0.1%) and then blocked in blocking buffer (5% normal goat serum in PBS-Triton X-100) for 30 minutes at room temperature. Samples were incubated at room temperature with rabbit anti–PLP (Rogers and Rogers, 2008) diluted at 1:3,000 in blocking buffer followed by three washes with PBS-Triton (0.1%). Cells were next incubated with Rhodamine red-X goat anti-rabbit secondary antibody (Jackson ImmunoResearch, diluted 1:1,500) and Hoechst 33342 (Life Technologies, 3.2 μM final concentration) in blocking buffer, followed by three washes with PBS-Triton X-100 (0.1%). Samples were mounted in Vectashield (Vector Laboratories). Super-resolution microscopy was performed using a Zeiss ELYRA S1 (SR-SIM) microscope equipped with an AXIO Observer Z1 inverted microscope stand with transmitted (HAL), UV (HBO) and solid-state (405/488/561 nm) laser illumination sources, a 100× objective (NA 1.4), 3 rotations, and an EM-CCD camera (Andor iXon). Images were acquired with ZEN 2011 software.

### Constructs

Full-length cDNAs of *Drosophila* Ana2, Plk4, and Sas4 were subcloned into a pMT vector containing in-frame coding sequences for EGFP, V5, or myc under control of the inducible metallothionein promoter. PCR-based site-directed mutagenesis with Phusion DNA polymerase (ThermoFisher) was used to generate the Ana2, Sas4 and Plk4 point mutants. Transgenic constructs were induced to express ∼24 hrs after transfection by the addition of 0.1-1 mM copper sulfate to the culture medium.

### Immunoprecipitation assays

GFP-binding protein (GBP) (Rothbauer et al., 2008) was fused to the Fc domain of human IgG (pIg-Tail) (R&D Systems), tagged with His_6_ in pET28a (EMD Biosciences), expressed in *E. coli* and purified on HisPur Cobalt resin (Fisher) according to manufacturer’s instructions (Buster et al., 2013). Purified GBP or mouse anti-V5 monoclonal antibody (Life Technologies) was bound to magnetic Dyna Beads (Fisher), and then cross-linked to the resin by incubating with 20 mM dimethyl pimelimidate dihydrochloride in PBS, pH 8.3, 2 hours, 22°C, and then quenching the coupling reaction by incubating with 0.2 M ethanolamine, pH 8.3, 1 hour, 22°C. Antibody-coated beads were washed three times with PBS-Tween 20 (0.02%), then equilibrated in 1.0 ml of cell lysis buffer (CLB; 50 mM Tris, pH 7.2, 125 mM NaCl, 2 mM DTT, 0.1% Triton X-100, 1x protease inhibitor cocktail [Roche], 0.1 mM PMSF, 1x phosphatase inhibitor cocktail set 1 [Sigma]). For replacement experiments, cells were transfected with dsRNA on day 0, followed by concurrent transfection with 1-3 µg plasmid (encoding the exogenous construct) and 20 µg dsRNA on day 4. Expression of exogenous protein was induced with copper sulfate on day 5, and samples were collected on day 6. Transfected cells expressing recombinant proteins were lysed in CLB and the lysates clarified by centrifugation at 16,100 × g for 5 minutes, 4°C. Transfected cells for mass spectrometry were not pre-cleared. 0.5-1% of the inputs were used for immunoblots. GBP-coated beads were rocked with lysates for 30 min, 4°C (or with anti-V5-coated beads for 10 min, 22°C), then washed four times with 1 ml CLB, and boiled in Laemmli sample buffer.

### Protein purification and pre-phosphorylation

GST-Ana2 FL (full-length) and 1-60 (amino acids 1-60) were purified as previously described (McLamarrah et al., 2018). Briefly, GST-tagged Ana2 construct was bacterially expressed and then purified from clarified bacterial lysate using glutathione resin (Fisher) following manufacturer’s instructions. Heat shock protein was removed by equilibrating resin-bound GST-Ana2 in 10 column volumes of HSP Buffer (50 mM Tris, pH 7.5, 50 mM KCl, 20 mM MgCl_2_) followed by washing with 20 column volumes of HSP Buffer containing 5mM ATP (Sigma).

GST tag was removed from GST-Ana2 by incubating the resin-bound protein with PreScission Protease (GE Healthcare) overnight, 4°C. Protease-cleaved protein was recovered in the column flow-through, and the material was then passed through new glutathione resin to remove any remaining GST-tagged protease and uncut GST-tagged Ana2. Samples were analyzed by SDS-PAGE.

His_6_-Thr-Sas4-606 (amino acids 606-901) and His_6_-FLAG-Plk4 (amino acids 1-317) were purified on HisPur Cobalt resin (Fisher) according to manufacturer’s instructions. His_6_-Thr-Sas4-606 was eluted from the resin with 200 mM imidazole in PBS, 1mM DTT followed by concentration and buffer exchange (into PBS, 1mM DTT) using Amicon Ultra spin concentrators (EMD Millipore). His_6_-FLAG-Plk4 1-317 was eluted similarly and exchanged into reaction buffer (RXN; 40 mM Na HEPES, pH 7.3, 150 mM NaCl, 5mM MgCl_2_, 1mM DTT, 10% [by volume] glycerol).

GST-Ana2 (FL and 1-60) was phosphorylated prior to cleavage of the GST-tag as follows. GST-Ana2 immobilized on glutathione resin was exchanged into RXN buffer and incubated with 20 µM purified His_6_-FLAG-Plk4 1-317 and 100 µM ATP for 1 hour at room temperature while shaking. To remove Plk4, the resin (with immobilized Ana2) was washed with 10 column volumes (CV) of wash buffer (PBS, 1 mM EGTA, 1 mM PMSF, 1 mM DTT, 1 mM soybean trypsin inhibitor [SBTI]), then 10 CV of wash buffer containing an additional 500 mM NaCl, followed by another 20 CV of wash buffer. The GST-tag was cleaved overnight as described above, releasing purified tagless Ana2.

### pExpress-Dual screen for isolating soluble protein complexes

We developed a four-step high-throughput method to screen for stable, soluble binary protein complexes from a library of constructs via co-expression in *E. coli*.

#### Step 1: Design and cloning for pExpress-Dual screen

pExpress-Dual plasmid was constructed by PCR amplifying a cassette containing a TEV cleavage site (ENLYFQS) followed by the linker GGSGGGSGGGSG and EGFP coding sequence (without a start codon) with two stop codons (TAG-TAA), which was then cloned into pST39 between the T7 promoter and the T7 terminator using primers TEV-linker F: 5’-CTCACTATAGGGAGACCACAACGGTTTCCCGAAAACCTGTACTTCCAATCCggtggttct-3’ and GFP-Stop R: 5’-TCCTTTCGGGCTTTGTTAGCAGCCGGATCTTTACTACTTGTACAGCTCGTCCATGCCGAG-3’. To assemble the dual expression library, pExpress-Dual was amplified as two pieces: (*1*) pExpress A using primers L3F: 30 bp of gene-specific sequence for each gene/fragment and 5’-GAAAACCTGTA CTTCCAATCCggtggttct-3’ and L4R: 5’-TTTATCAGCAATAAACCAGCCAGCAGGAAGTGCCGA-3’, and (*2*) pExpress B using primers L4F: 5’-AAAGTTGCAGGACCACTACTG CGTTCGGCA-3’ and L1R: 5’-ATTTCTCCTCTTTAATTCTAGGTACCCGGGGGGAAACCGTTGTGGTCTCCCTATAGTGAG-3’. Primers for gene A cloning include L1F: 5’-CCC GGGTACCTAGAATTAAAGAGGAGAAATTAAGCATGCACCACCATCACCATCAT-3’ and L2R: 5’-TATTCGACTATAACTGGCCGTCGTTTTACATTA-3’ followed by 30 bp of gene-specific sequence. Primers for gene A cloning include L2F: 5’-TGTAAAACGACGGCCAGTTATAGTCGAATAAAACAAGTGGCTAAGGAGGTTGT-3’ and L3R: 5’-agaaccacc GGATTGGAAGTACAGGTTTTC-3’. PCR products ≥1 kb were mixed at equimolar ratios and at 5x molar ratios for fragments <1 kb. 30 ng of pExpress B and 15 μl of master mix were added for each 5 μl of DNA mixture and incubated for two hours at 50°C and transformed (Gibson, 2011).

#### Step 2: Bacterial protein expression. (A) Transformation

1 µl of the plasmid library reaction mixture was added to 50 µl BL21(DE3) pLysS competent *E. coli*, incubated for 20 minutes on ice, and heat-shocked for 1 minute at 37 °C. 300 µl of LB was added to the mixture, placed in a 250 rpm shaker for 1 hour at 37 °C, plated on 3 agar plates supplemented with 100 µg/m ampicillin, and incubated overnight at 37 °C. *(B) Colony expansion*. 96 random colonies were inoculated into 96-well plates containing 100 µg/ml ampicillin and replicate plated on gridded LB amp plates. *(C) Bacterial growth and induction of protein expression*. 1 ml of TB medium with ampicillin and 1% (v/v) glucose was added to each well, plates were sealed with Parafilm® and placed in a 250 rpm shaker at 37 °C overnight. 100 µl of culture was diluted into a new 96-well plate containing 900 µl of supplemented TB medium per well, sealed and incubated at 250 rpm at 37°C until cultures reached an OD_600_ = 1.0. Expression was induced by adding 0.5 mM IPTG per well and plates were incubated at 250 rpm at 18°C overnight. Cells were harvested by centrifugation at 2000xg at 4 °C for 15 minutes, supernatants decanted, and plates stored at −80 °C. *(D) Purification of soluble protein complexes* 150 µl of lysis buffer was added to each well, plates were placed in a shaker at 30°C for 30 minutes, and then incubated on ice for 5 min. 50 µl of Benzonase® buffer was added to each well and agitated at 4°C for 15 minutes. Lysates were clarified by centrifuging plates at 2000xg for 15 minutes at 4°C. 50 µL of equilibration-binding buffer was added to each well, transferred to a pre-equilibrated HisPur™ Ni-NTA Spin Plates, sealed and agitated at 4°C for 30-60 minutes. Plates were placed on a vacuum manifold and filtered using ∼4 inches Hg of pressure, washed 3x with 200 µl of wash buffer and resin resuspended in 100 µl of elution buffer. Plates were then sealed and incubate with gentle agitation at 4°C for 15 minutes. Holes were punched on either side of a clean 96-well collection plate, the filter plate and the collection plate were sealed, and eluate collected by vacuum pressure.

#### Step 3: Primary Screening

2 µl of eluate from each well was transferred onto 2 nitrocellulose membranes. For GFP-immunoblotting protein detection, membranes were blocked with 5% milk in TBS containing 0.05% Tween-20 (TBST) for 1 hour at room temperature. For detection of His_6_-tagged protein, membrane was blocked with 2.5% BSA in PBS containing 0.3% Triton X-100 (PBST) for 1 hour at room temperature.

#### Step 4: Secondary Screening

False positives from the primary screen was identified by: *(A)* Single protein insertions that were double-tagged with His_6_ on the N-term and GFP on the C-terminus were identified by PCR using T7 and M13F (-21) reverse primers that recognize flanking regions of gene A insertion sites. Any colony not producing PCR product was marked as a dual-tagged false positive. Following this step, 62 colonies remained. *(B)* Duplication were identified by sequence analysis, yielding 48 candidate dual expression vectors. *(C)* Tandem pulldown/IP for His_6_ and GFP. For His_6_ IPs, 25ml overnight cultures were pelleted and resuspended in 1 ml of lysis buffer, incubated on ice for 5 minutes and sonicated. Lysates were then precleared by centrifugation at 2000xg for 15 minutes at 4°C. 25 µl of equilibrated His-Tag Dynabeads (total bed volume) were added to supernatant and rocked for 15-30 minutes at 4 °C. Beads were retrieved with a magnet, gently washed 3x with 300 µl wash buffer and resuspended in 150 µl of elution buffer for 10 minutes at 4 °C. For GFP IPs, 25 µl of Protein-A Dynabeads (total bed volume) were resuspended in 1 ml PBS + 0.01% Tween-20 with 0.5 µl of ab290 anti-GFP antibody and rocked for 30-60 minutes at 4°C. Proteins were eluted by resuspending beads in 25-50 µl of SDS-PAGE loading buffer (2X) and boiling for 8 min.

#### Buffer recipes for pExpress-Dual screen

Luria Broth (LB) medium (1.0% tryptone, 0.5% yeast extract, 1.0% NaCl); Terrific Broth (TB) medium (1.2% tryptone, 2.4% yeast extract, 0.5% (v/v) glycerol, 0.17 M KH2PO_4_, 0.72 M K_2_HPO_4_). Lysis buffer (B-PER™ Bacterial Protein Extraction Reagent, 150 mM NaCl, 10 mM 2-mercaptoethanol, 0.1 mM EDTA, 0.2 mg/ml Lysozyme, 1 mM PMSF, cOmplete™ EDTA-free Protease Inhibitor Cocktail). Benzonase buffer (50 mM Tris-HCl, pH 7.4, 1.2 mM MgCl2, 0.1 U/uL Benzonase®). Equilibration/Binding buffer (50 mM Tris-HCl, pH 7.4, 150 mM NaCl, 5 mM imidazole, 10 mM 2-mercaptoethanol, 1 mM PMSF, cOmplete™ EDTA-free Protease Inhibitor Cocktail). Wash buffer (50 mM Tris-HCl, pH 7.4, 150 mM NaCl, 10 mM imidazole, 10 mM 2-mercaptoethanol, 1 mM PMSF, cOmplete™ EDTA-free Protease Inhibitor Cocktail). Elution buffer (50 mM Tris-HCl, pH 7.4, 150 mM NaCl, 300 mM imidazole, 10 mM 2-mercaptoethanol).

### Bio-layer interferometry

Bio-layer interferometry data was collected on a Pall ForteBio Octet RED384 system using 96-well plates from Grenier (cat no: 655209). Kinetic assays were performed by first equilibrating Ni-NTA Octet biosensors (ForteBio Lot # 1810232) in kinetic buffer (PBS, 0.01% BSA, 0.002% Tween 20) during a baseline step for 60 s followed by loading with 8 µg/mL His_6_-Thr-Sas4-606 in PBS, 1 mM DTT for 120 s. Sas4-coated biosensors were next equilibrated in sample buffer (PBS, 0.01% CHAPS, 1% BSA, 10 mM imidazole) for 180 s in a second baseline step. Sas4 captured biosensors were then submerged in wells for 120 s containing different concentrations of either Ana2 FL or 1-60, or phosphorylated Ana2 FL or 1-60. This association step was followed by a dissociation step for 300 s in sample buffer. Ni-NTA biosensors were regenerated using a 5 s regeneration step in 500 mM imidazole in PBS followed by a 5 s neutralization step in kinetic buffer. This process was repeated 5 times for complete regeneration of biosensors. The binding sensorgrams were collected using the 8-channel detection mode on the Octet RED384. Association and dissociation constants were calculated with Pall ForteBio Data Analysis 11.0 software. Binding sensorgrams were globally fit in a 1:1 binding model following subtraction of a single reference set. Data was aligned in the y-axis to the average of the baseline step with an inter-step correction aligned to the dissociation step and filtered using Savitzky-Golay filtering. Graphs were generated using GraphPad Prism.

### In silico structural analysis

Full-length Ana2 WT or Ana2 mutant sequence files were uploaded to the I-TASSER website (**I**terative **T**hreading **ASSE**mbly **R**efinement; https://zhanglab.ccmb.med.umich.edu/I-TASSER) for protein structure and function prediction (Zhang, 2008; Roy et al., 2010, Yang et al., 2016). Using I-TASSER, five models were predicted for each construct, and an optimal model was selected for each construct based on C-score and known structural features. The normalized B-factor (B-factor profile or BFP) is predicted by the I-TASSER algorithm using a template-based and profile-based prediction (Yang et al., 2016).

### Mass spectrometry

S2 cells expressing transgenic Ana2 were solubilized in cell lysis buffer (CLB) and tagged Ana2 was purified either by immunoprecipitation (using anti-V5) or pull-down (using GFP binding protein, GBP). Samples were further resolved by SDS-PAGE, and selected regions of the Coomassie-stained gels were collected. After destaining, gel pieces were reduced (10 mM DTT in 25 mM NH_4_HCO_3_, 55°C, 60 min), alkylated (55 mM iodoacetamide in 25 mM NH_4_HCO_3_, 45 min, in dark at room temperature), and proteolyzed (1.5 µg chymotrypsin, 0.6 µg trypsin in 50 mM NH_4_HCO_3_, 1 mM CaCl_2_, overnight at room temperature, then 2 hrs at 37°C). Peptides were extracted from the gel pieces (using a series of 20 min incubations at 37°C in 1% trifluoroacetic acid [TFA] and 25-99% acetonitrile), dried and resuspended in 0.1% TFA, and then desalted with NuTip carbon micro-cartridges (Glygen). Processed samples were stored at −20°C until analyzed.

LC-MS/MS analysis was carried out using a Q Exactive Plus mass spectrometer (Thermo Fisher Scientific) equipped with a nanoESI source. The nanoLC was a Dionex Ultimate 3000 RSLCnano System (Thermo Scientific). Peptides were loaded onto an Acclaim Pepmap 100 trap column (75 µm ID × 25 cm, Thermo Scientific) for 10 min at 3 µLs/min then eluted onto an Acclaim PepMap RSLC analytical column (75 µm ID × 25 cm, Thermo Scientific) using a 5-20% gradient of solvent B (acetonitrile, 0.1% formic acid) over 122 min, 20-50% solvent B over 10 min, 50-95% of solvent B over 10 min, 95% hold of solvent B for 10 min, and finally a return to 5% solvent B in 1min and another 10 min hold of 5% solvent B. Solvent A consisted of water and 0.1% formic acid. Flow rate on the analytical column was 300 nL/min. Spectral data were acquired using a Data Dependent scanning by Xcalibur software version 4.0.27.19 (Andon et al., 2002), using a survey scan at 70,000 resolution scanning mass/charge (m/z) 353-1550 at an automatic gain control (AGC) target of 1×10^5^ and a maximum injection time (IT) of 65 msec, followed by higher-energy collisional dissociation (HCD) tandem mass spectrometry (MS/MS) at 27nce (normalized collision energy) of the 10 most intense ions at a resolution of 17,500, an isolation width of 1.5 m/z, an AGC of 1×10^5^, and a maximum IT of 65 msec. Dynamic exclusion was set to place any selected m/z on an exclusion list for 20sec after a single MS/MS. Ions of charge state +1, 7, 8, >8 and unassigned were excluded from MS/MS, as were isotopes.

Tandem mass spectra were searched against a *Drosophila melanogaster* proteome Uniprot database to which common contaminant protein (e.g., chymotrypsin, trypsin, keratins; obtained at ftp://ftp.thegpm.org/fasta/cRAP) and the transfected construct sequences were appended. All MS/MS spectra were searched using Thermo Proteome Discoverer v2.2.0388 (Thermo Fisher Scientific) considering fully proteolyzed peptides with up to 2 missed cleavage sites. Variable modifications considered during the search included methionine oxidation (15.995 Da), cysteine carbamidomethylation (57.021 Da), and phosphorylation (79.99 Da) of serine, threonine and tyrosine. Proteins were identified at 99% confidence with XCorr score cut-offs (Qian et al., 2005) as determined by a reversed database search. The protein and peptide identification results were also visualized with Scaffold Q+S version 4.8.7 (Proteome Software), which relies on the Sequest search engine results and uses Bayesian statistics to reliably identify more spectra (Keller et al., 2002). Assignment of the phosphorylation sites listed in the Supplemental Table was performed with Scaffold PTM (Proteome Software); S38, S63, and S150 of Ana2 expressed in S2 cells (co-expressing transgenic Sas4) were identified as phospho-sites with high confidence (Ascore > 18). However, the site assignments of the S150-containing phosphopeptides obtained from the control samples were low confidence (i.e., Ascore < 18), so the actual position of the single phosphate moiety within these particular phosphopeptides is unknown. We report the phosphopeptide sequences with the largest Ascores.

## Online supplemental material

Fig. S1 shows that replacement of Sas4 with an Ana2-binding mutant (ABM) prevents hyperphosphorylation of endogenous Ana2. Additionally, the Sas4 G-box is sufficient to promote Ana2 hyperphosphorylation but not when containing the ABM. Proteins that were screened for binary complex formation and shown. Fig. S2 show a flowchart of the binary protein screen. Fig. S3 and S4 show workflows of the primary and secondary screens, respectively. Fig. S5 shows that phosphorylation of the STAN domain is not responsible Ana’s electrophoretic shift but that phospho-S38 is necessary and sufficient for this shift. Fig. S6 shows the purified of proteins used in the bio-layer interferometry assay and data from those experiments. Fig. S7 shows protein structure predicts for Ana2 variants. Table S1 lists primers used in constructing the pExpress vectors for the protein binding screen. Table S2 lists Ana2 phospho-sites identified by tandem MS in this study and

## Acknowledgements

We thank S. Smith for help confirming Plk4-Sas4 interactions and A. Kelly for help with screen design. This work was supported by the Division of Intramural Research at the NIH/NHLBI (1ZIAHL006104) to N.M.R. G.C.R. is grateful for support from NCI P30 CA23074 and NIGMS R01 GM110166 and GM126035NIH/NIGMS R01GFM110166.

Authors declare no competing financial interests.

## Supplemental materials

**Figure S1.**
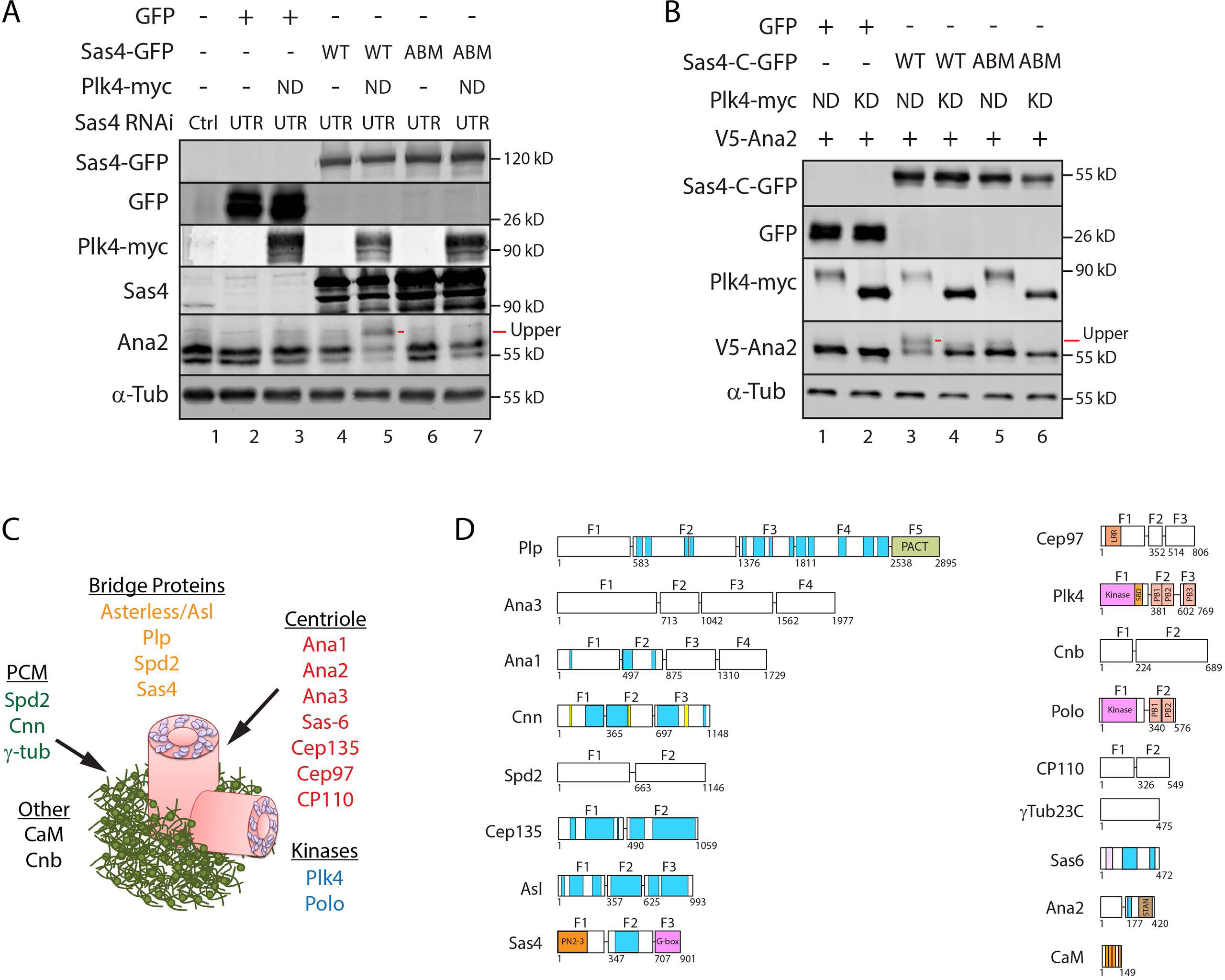
Expression of Ana2-binding mutant (ABM) Sas4 prevents Ana2 hyperphosphorylation. **(A)** Endogenous Ana2 does not shift to the slower migrating hyperphosphorylated species when endogenous Sas4 is replaced with Sas4-ABM. S2 cells were depleted of endogenous Sas4 by targeting its untranslated region (UTR) for 12 d. On days 4 and 8, cells were transfected with the indicated constructs and induced to express the next day for the duration of the experiment by the addition of 1mM CuSO_4_. Proteins were detected by immunoblotting cell lysates with anti-GFP, myc, Sas4, Ana2 and α-tubulin. **B** Expression of Sas4-C (G-box) harboring the ABM prevents the electrophoretic shift in transgenic Ana2. S2 cells were treated as in (**A**). Proteins were detected by immunoblotting cell lysates with anti-GFP, myc, V5, and α-tubulin. **C** Cartoon shows the 17 centrosome proteins that were selected for binary complex solubility screening and their positions within the centrosome. **D** Schematic of the proteins tested in the screen. Numbers show amino acids, blue regions predict coiled-coils, additional structural/functional domains are indicated, and horizontal lines show the locations where proteins were subdivided into smaller testable fragments (F1-5). Also described in Galletta et al. (2016).

**Figure S2.**
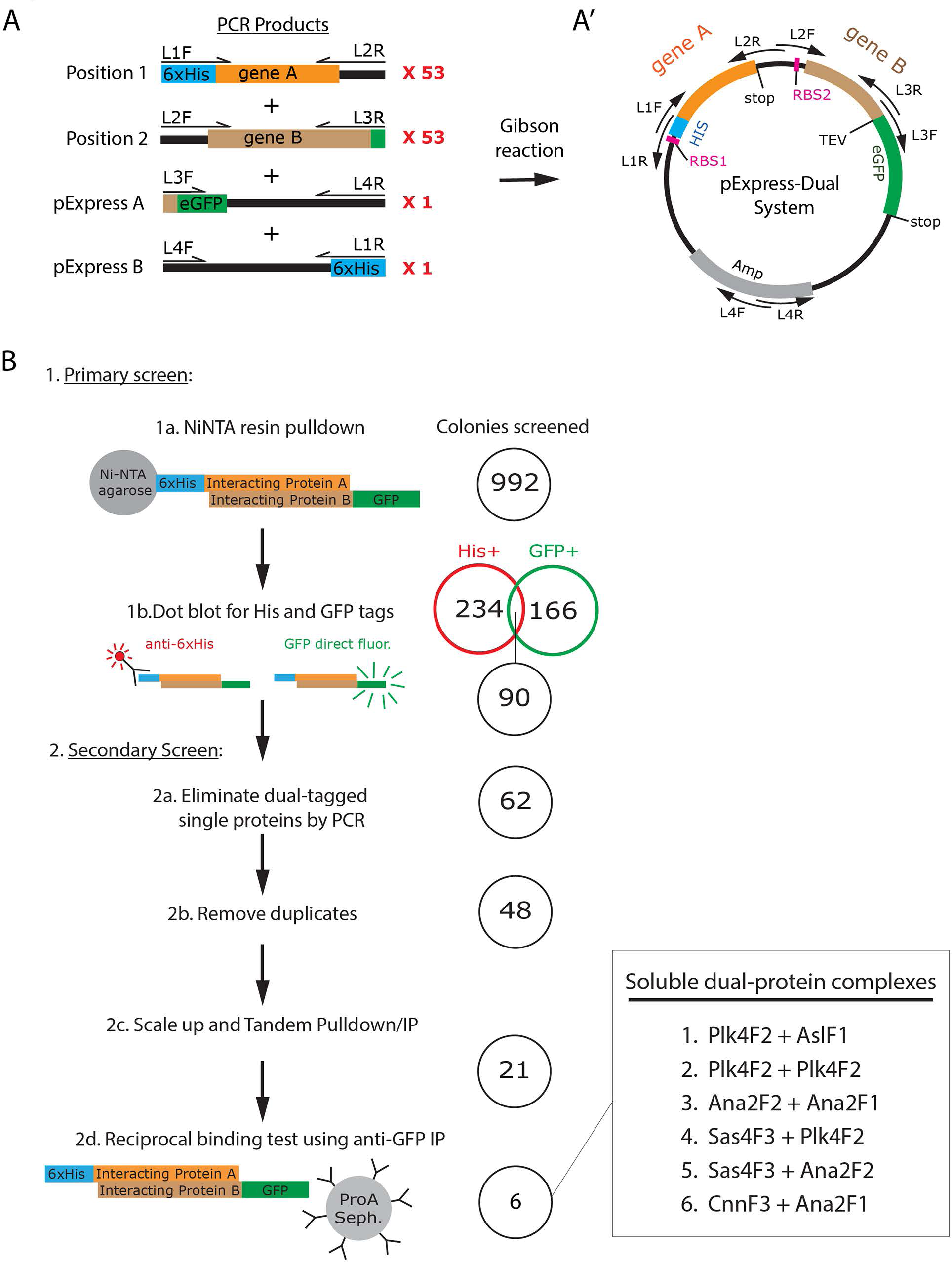
A dual-expression screen of *Drosophila* centrosome proteins identifies 6 soluble binary protein complexes. **(A)** PCR products used for random plasmid assembly. Each gene was amplified as full-length and/or sub-fragments using primers listed in Table S2. L1F and L2R primers were used to generate 53 unique PCR products that could occupy the ‘gene A’ position. L2F and L3R primers were used to generate 53 unique PCR products that could occupy the ‘gene B’ position. pExpress A and B PCR fragments were used to assemble the plasmid backbone. **(A’)** Schematic of the final plasmid indicating the positions of the His_6_-geneA and geneB-GFP, primers, and the Ribosome Binding Sites (RBS1, RBS2). **(B)** Summary of screen and results. A total of 992 colonies were cultured and processed as described. (1a and b) The was conducted in two steps. Primary screening used Ni-NTA pulldowns followed by immuno-dot blots to detect His and GFP-tagged protein, resulting in 90 binary complexes. The secondary screen consisted of 4 steps: (2a) Using PCR to elimination individual proteins that were dual tagged with both His and GFP, (2b) the removal of duplicate plasmids, (2c) scaling up the culture for tandem pulldown/IP using Ni-NTA then GFP antibodies, and finally (2d) repeating the 21 positive hits with a reciprocal IP using anti-GFP. This resulted in the isolation of the 6 protein complexes listed. Complexes 1, 2 and 3 were previously identified by other studies and serve as a strong proof of concept (Dhindzhev et al., 2010; Slevin et al., 2012; McLamarrah et al., 2018). Interactions 4 and 5 are the subject of this study. Interaction 6 is a novel interaction.

**Figure S3.**
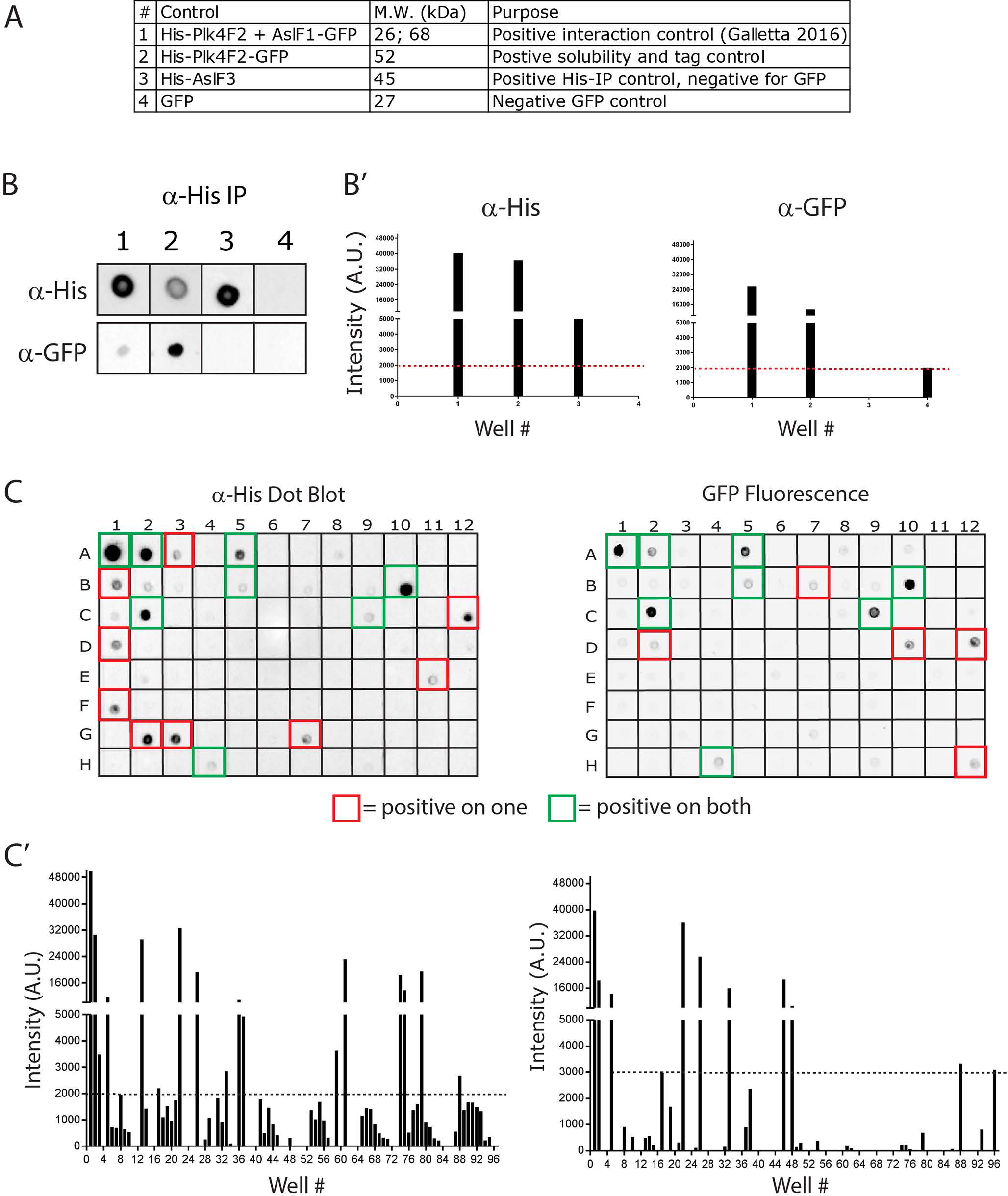
Workflow of the primary screen. **(A)** Four controls were used in each 96-well screening plate. Description of the controls, expected molecular weights, and the purpose of each are indicated. **B** Immuno-dot blots of controls indicated in A. Elutes from the HisPur™ Ni-NTA Spin Plates were blotted for His and GFP. (**B’**) Signal from anti-His HRP-conjugated antibody and from GFP were measured and plotted using ChemiDoc MP Imaging System (Bio-Rad). Intensity plots (A.U. = arbitrary units) were used to determine positive hits for both His and GFP. **C** A representative example of a dot blot (Left for His, Right for GFP) from an entire 96-well plate with controls in positions 1-4. Hits are indicated with colored boxes: Single positive (red) and double positive (green) for both His and GFP are indicated. **(C’)** Intensity measurements of the corresponding the dot blots in C.

**Figure S4.**
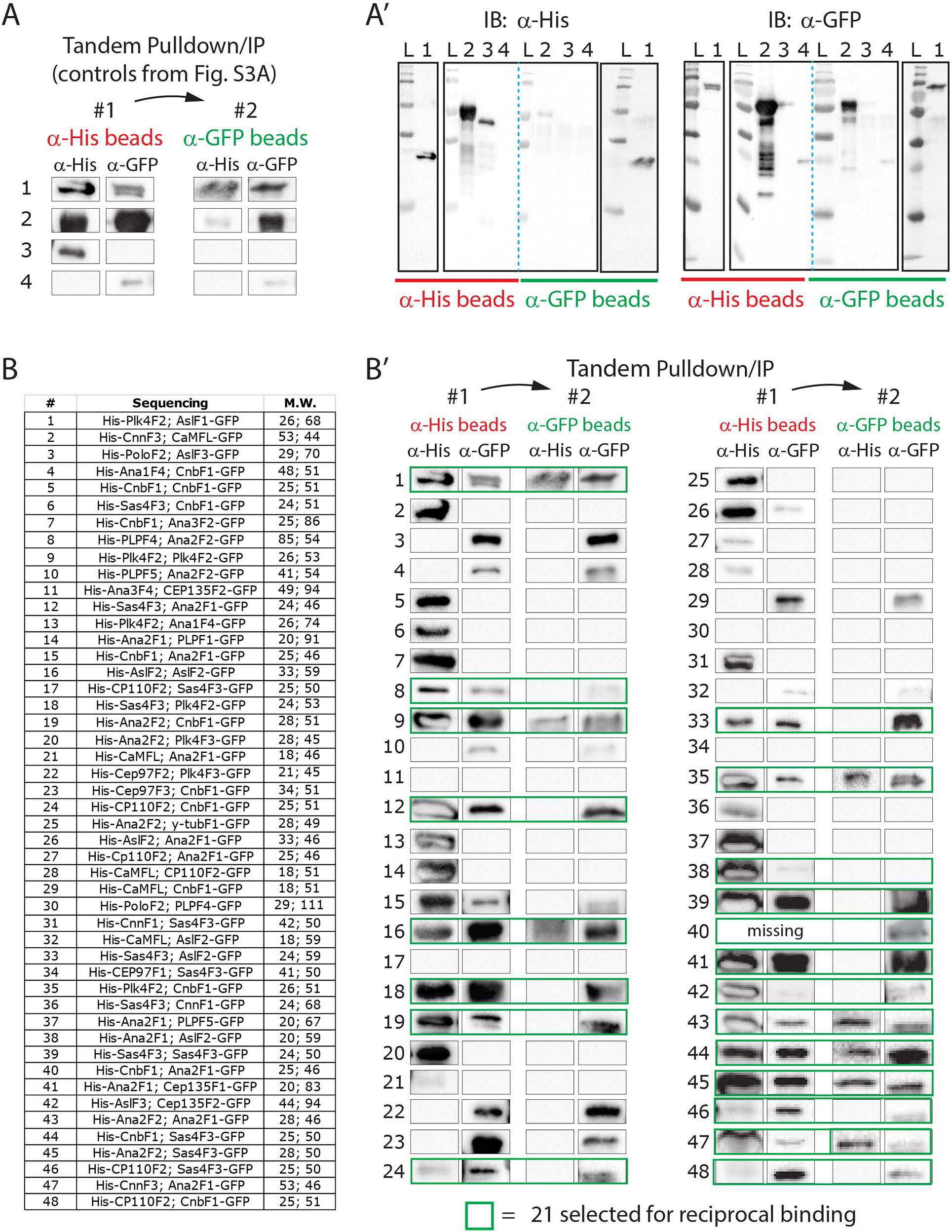
Workflow of the secondary screen. **(A)** The identical four controls (Fig. S3 A) were used for secondary screening which began with scaled-up (25 ml) cultures of the 48 positive hits (Figure S2 B, step 2b). Lysates were processed through a tandem pulldown/IP protocol using His-tag Dynabeads and then anti-GFP-conjugated beads. Note, control #4 immunoblots indicates that soluble GFP binds His-tag Dynabeads at low levels. **(A’)** Immunoblots of the original controls that were used to generate panel A. **(B)** List of all 48 hits that were screened by tandem pulldown/IP indicating the genes that occupied Positions 1 and 2 in the pExpress-Dual plasmid as well as their molecular weights. **(B’)** Immunoblots show results of the tandem pulldown/IP for all 48 hits. Note, immunoblots of the His-tag pulldown for #40 is missing. All 21 green boxed blots were selected for the final stage of screening: the reciprocal binding test performed by anti-GFP IP (Fig. S2 B, step 2b).

**Figure S5.**
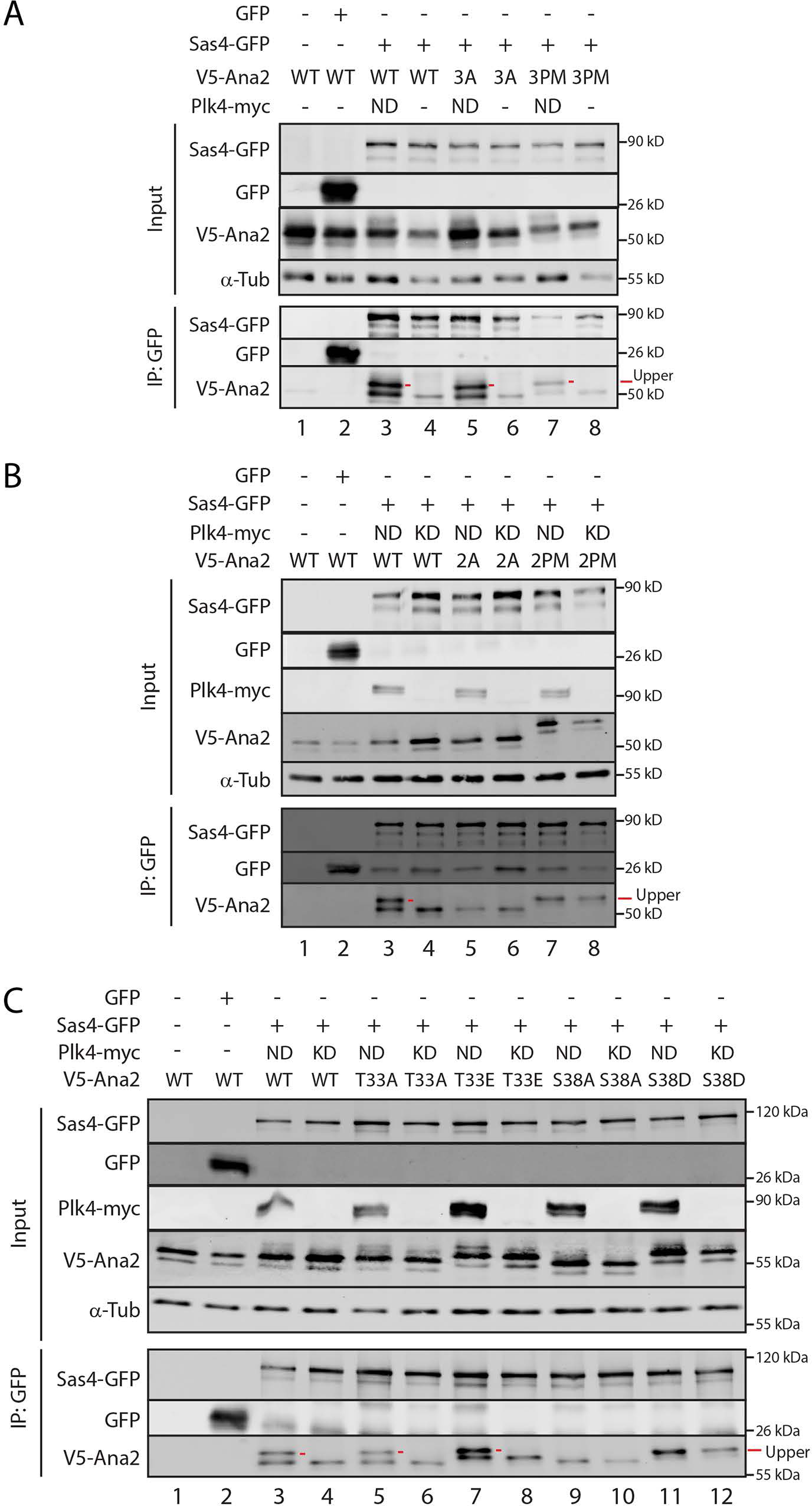
S38 is the phospho-residue in Ana2 responsible for its electrophoretic shift. **(A)** Phospho-mutants in the STAN domain (3A or 3PM) of Ana2 do not affect the ability of Sas4 and Plk4 to hyperphosphorylate Ana2. S2 cells were co-transfected with the indicated constructs and the next day induced to express for 24 hours by addition of 1 mM CuSO_4_. Anti-GFP IPs were then prepared from lysates, and Western blots of the inputs and IPs probed for GFP, V5, and α-tubulin. **(B)** Double phospho-mutation of N-terminal residues T33 and S38 (2A or 2PM) alter the electrophoretic shift of Ana2. S2 cells were treated as in (**A**). Anti-GFP IPs were then prepared from lysates, and Western blots of the inputs and IPs probed for GFP, myc, V5, and α-tubulin. **(C)** Phospho-mutations in S38 but not T33 alter the electrophoretic shift of Ana2. S2 cells were treated as in (**A**). Anti-GFP IPs were then prepared from lysates, and Western blots of the inputs and IPs probed for GFP, myc, V5, and α-tubulin.

**Figure S6.**
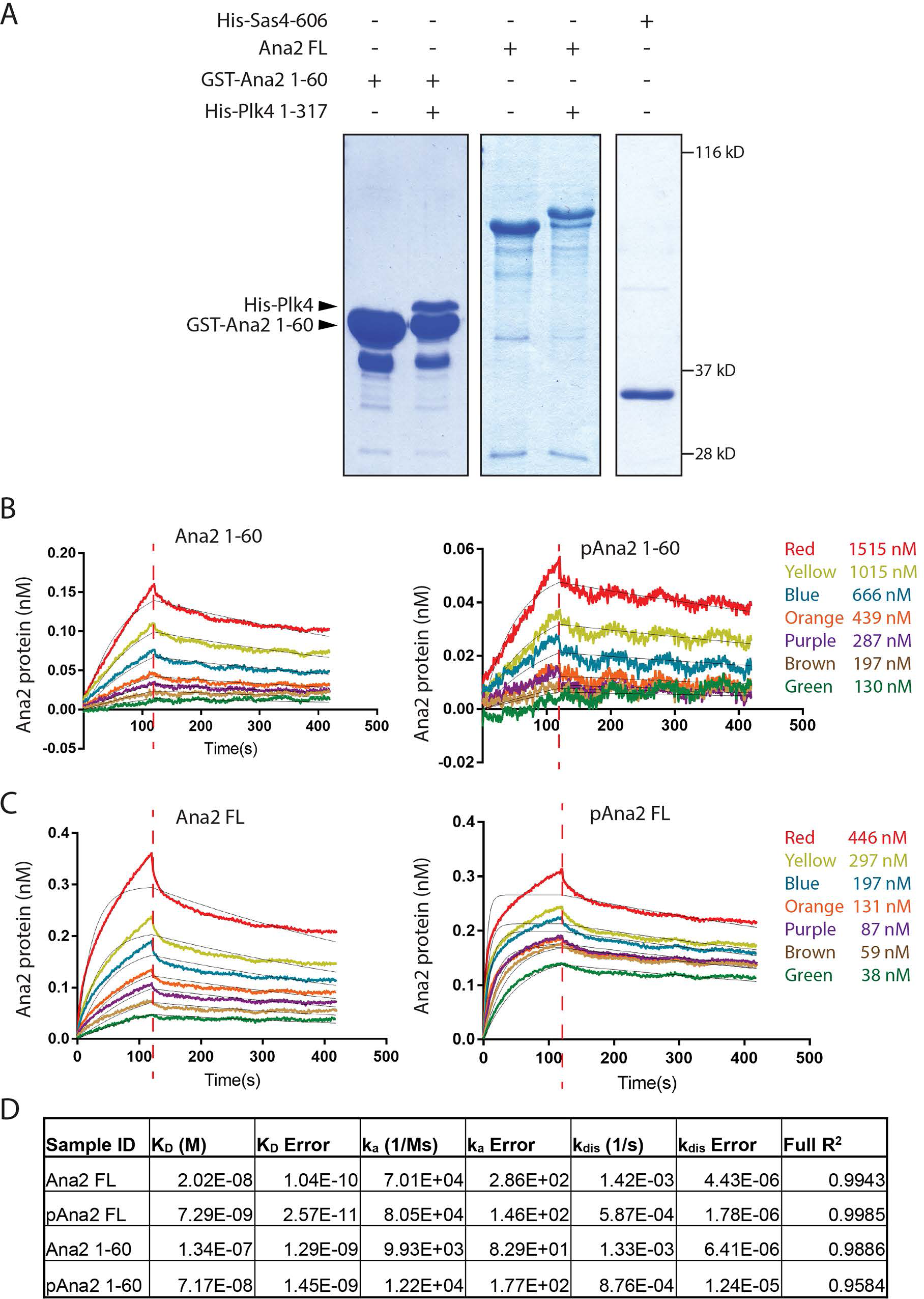
Full-length phospho-Ana2 binds tighter to the Sas4 G-box in vitro. **(A)** Coomassie-stained protein gels showing the purity of the proteins used in the in vitro binding kinetics study. Proteins were purified as GST or His_6_-tagged proteins. The GST tags on full-length (FL) and 1-60 Ana2 were proteolytically removed. For some treatments, Ana2 was phosphorylated with purified Plk4 kinase domain (amino acids 1-317), repurified, and then used for the binding experiments. **(B and C)** Graphs show dissociation/binding kinetics for untreated and phospho-Ana2 proteins with the Sas4 G-box using the Octet RED384 system. The Ana2 proteins were used at the indicated range of concentrations and G-box protein was used at a constant 216 nM. Dashed red lines mark the start of the dissociation step. **(D)** Measured binding parameters between purified Sas4 G-Box and Ana2 proteins (untreated and phosphorylated) full-length (FL) and the 1-60 fragment from bio-layer interferometry. A single data set from each binding experiment is shown.

**Figure S7.**
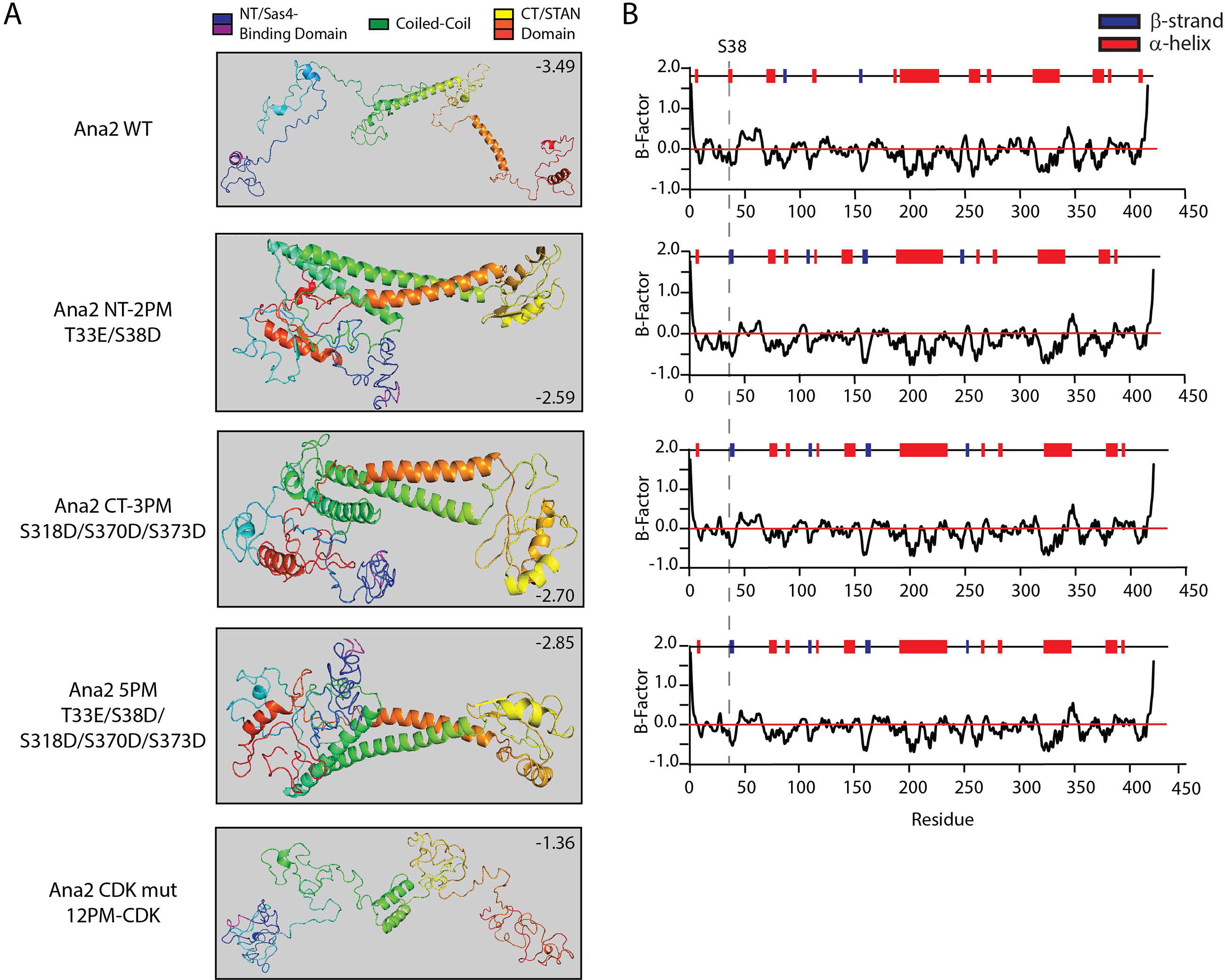
Phosphomimetic substitutions in the N- and C-terminal domains of Ana2 predict a folded conformation with close positioning of the terminal domains. **(A)** I-TASSER protein structure predictions of wild-type Ana2 and several phosphomimetic Ana2 mutants. C-scores are shown. Previously, we showed that Ana2 is first phosphorylated on N-terminal (NT) residues T33/S38, then rapidly phosphorylated on C-terminal (CT) residues S318/S370/S373 (McLamarrah et al., 2018). Phosphomimetic (PM) substitutions in either the Ana2 NT (2PM) or CT (3PM) promotes in trans binding between the two domains; maximal binding occurred between NT-2PM and CT-3PM (McLamarrah et al., 2018). Notably, NT-2PM, CT-3PM or the full 5PM mutant predict a folded conformation with close positioning of the termini. Note, that the 12PM CDK ([S/T]P) Ana2 mutant is not predicted to generate a folded conformation. **(B)** I-TASSER predictions of secondary structure within the corresponding Ana2 proteins in (**A**). The position of S38 is indicated (dashed line) in a short segment that changes from a α-helix to a β-strand in the phosphomimetic Ana2 mutant variants.

**Table S1.**
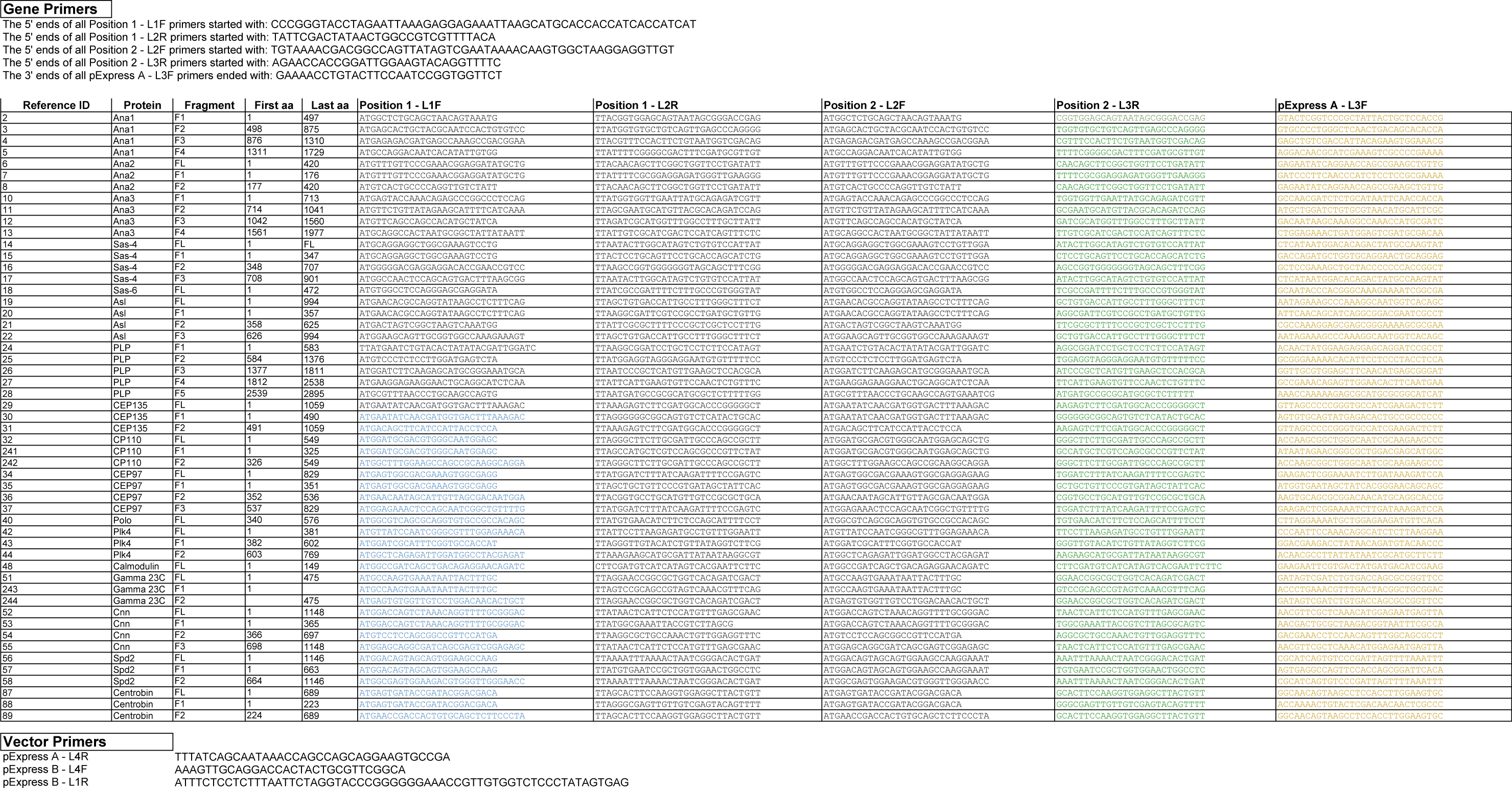
List of gene primers used for the pExpress-Dual screen for isolating soluble protein complexes.

**Table S2.** Sas4 catalyzes Plk4-mediated phosphorylation of Ana2 S38, producing a hyperphosphorylated form of Ana2 as determined by tandem mass spectrometry of Ana2 from S2 cells. **(A)** S2 cells were co-transfected with plasmids encoding the inducible transgenic constructs GFP-Ana2 and non-degradable (ND) Plk4 (an active and degradation-resistant mutant of Plk4). In addition, these cells were transfected with either (1) a plasmid encoding Sas4-V5 to overexpress Sas4 (‘+ Sas4-V5 OE’), or (2) Sas4 dsRNA to deplete the endogenous Sas4 protein (‘+ Sas4 RNAi’). As described in Methods, GFP-Ana2 was IPed from S2 cell lysates and further resolved on SDS-PAGE. GFP-Ana2 bands were cut from the gel, processed to generate proteolytic fragments, and the mixtures of peptides analyzed by MS/MS to identify phosphorylated residues. The ScaffoldPTM program (Proteome Software) was used to identify the locations of phospho-modifications and to evaluate the reliability of each localization. Identified phospho-residues are indicated in the peptide sequences by small case font and underline. (A lowercase ‘m’ indicates methionine oxidation.) Note that, in two cases, the sites of modification within the phospho-peptides could not be reliably located (Ascores < 18); their phospho-residues and peptide sequences were selected by ScaffoldPTM. Spectrum counts were obtained by counting the number of spectra corresponding to peptides with the particular phospho-residue and also counting the total number of spectra of peptides containing the particular residue (regardless of phosphorylation state). Only NT phospho-modifications mentioned in the text are shown. **(B)** S2 cells were co-transfected with inducible expression plasmids encoding the transgenic constructs V5-Ana2 and Sas4-GFP. In addition, cells were transfected with either active ND-Plk4 or a kinase-dead (KD) mutant of Plk4. After IP of V5-Ana2 from cell lysates, samples were resolved by SDS-PAGE, and regions of the Coomassie-stained gel were collected as shown in Fig. 3 D. Samples were processed to generate extractable peptide fragments which were analyzed by MS/MS to identify phosphorylated residues (see Methods). Only the pS38 modification mentioned in Fig. 3 D is shown.

| Sample | Phospho Residue | Peptide Sequence | Phospho Localization Probability | Ascore | Peptide Score | Sequest: PeptideSp | Sequest: Peptide RankSp | Sequest: DeltaCN | Sequest: Xcorr | Scaffold: Peptide Probability | Actual Mass | Observed Mass | Charge | Delta AMU | Delta PPM | Spectrum Counts <sup>1</sup> |
| --- | --- | --- | --- | --- | --- | --- | --- | --- | --- | --- | --- | --- | --- | --- | --- | --- |
| **SECTION A** |  |  |  |  |  |  |  |  |  |  |  |  |  |  |  |  |
| **+ Sas4-V5 OE** | S38 | PSAAVPm<br>GHTNEIIGP<br>TVPEV<sub>s</sub>ILF | 100% | 40.54 | 152.97 | 177.3 | 1 | 0.1097 | 4.945 | 0.992 | 2671.307883 | 891.4432369 | 3 | 0.0108 | 4.03 | 5/98, (5%) |
|  | S63 | GQPPQD<br>PQmQPQ<br>QPNHAH<br>QES<sub>s</sub>PR | 99% | 18.44 | 118.89 | 151.4 | 1 | 0.008173 | 4.883 | 0.9895 | 2814.197457 | 704.5566402 | 4 | 0.00414 | 1.47 | 21/141,<br>(15%) |
|  | S150 | L<sub>s</sub>SSSNADV L | 98% | 20.98 | 79.5 | 300.6 | 1 | 0.3128 | 1.839 | 1 | 1071.448216 | 536.731384 | 2 | 0.000787 | -0.266 | 17/55,<br>(31%) |
| **+ Sas4 RNAi** | S38 | ND |  |  |  |  |  |  |  |  |  |  |  |  |  |  |
|  | S63 | ND |  |  |  |  |  |  |  |  |  |  |  |  |  |  |
|  | S150 | RL<sub>s</sub>SSSNADV L | 33% | 0 | 175.17 | 504.2 | 1 | 0.4541 | 3.384 | 1 | 1227.550877 | 614.7827145 | 2 | 0.00128 | 1.04 | 2/27, (7%) |
| **SECTION B** |  |  |  |  |  |  |  |  |  |  |  |  |  |  |  |  |
| **+ Plk4 KD** |  |  |  |  |  |  |  |  |  |  |  |  |  |  |  |  |
| Lower Region | S38 | ND |  |  |  |  |  |  |  |  |  |  |  |  |  |  |
| Upper Region | S38 | ND |  |  |  |  |  |  |  |  |  |  |  |  |  |  |
| **+ Plk4 ND** |  |  |  |  |  |  |  |  |  |  |  |  |  |  |  |  |
| Lower Region | S38 | ND |  |  |  |  |  |  |  |  |  |  |  |  |  |  |
| Upper Region | S38 | PSAAVP<br>mGHTN<br>EIIGPTV<br>PEV<sub>s</sub>ILF | 100% | 101.45 | 211.15 | 241 | 1 | 0.1556 | 4.72 | 0.9994 | 2671.3207 | 891.4475093 | 3 | 0.0236 | 8.83 | 3/8, (38%) |

ND, phospho modification not detected.
<sup>1</sup> # of phospho spectra / # of total spectra, (percent).

